# High admixture between forest an cultivated chestnut (*Castanea sativa* Mill.) in France

**DOI:** 10.1101/792259

**Authors:** Cathy Bouffartigue, Sandrine Debille, Olivier Fabreguette, Ana Ramos Cabrer, Santiago Pereira-Lorenzo, Timothée Flutre, Luc Harvengt

## Abstract

**Context:** Renewed interest in European chestnut in France is focused on finding locally adapted populations partially resistant to ink disease and identifying local landraces.

**Aims:** We genotyped trees to assess (i) the genetic diversity of wild and cultivated chestnut across most of its range in France, (ii) their genetic structure, notably in relation with the sampled regions, and (iii) relations with its neighbors in Spain and Italy.

**Methods:** A total of 693 trees in 16 sampling regions in France were genotyped at 24 SSRs, and 1,401 trees in 17 sampling regions at 13 SSRs.

**Results:** Genetic diversity was high in most sampling regions, with redundancy between them. No significant differentiation was found between wild and cultivated chestnut. A genetic structure analysis with no *a priori* information found a low, yet significant structure, and identified two clusters. One cluster gathers trees from south-east France and Corsica (RPP1) and another cluster gathers trees from all other sampled regions (RPP2). A substructure was detected at 13 SSRs suggesting differentiation within both RPP1 and RPP2. RPP1 was split between south-east France and Corsica. RPP2 was split between north-west France, Aveyron, Pyrenees and a last cluster gathering individuals from several other regions.

**Conclusion:** The genetic structure within and between our sampling regions is likely the result of natural events (recolonization after the last glaciation) and human activities (migration and exchanges). Notably, we provide evidence for a common origin of most French and Iberian chestnut trees, except those from south-east France that were associated with the Italian gene pool. This advances our knowledge of chestnut genetic diversity and structure, will benefit conservation and help our local partners’ valorization efforts.

## INTRODUCTION

Sweet chestnut (*Castanea sativa* Mill.) is an endemic, multi-purpose tree species cultivated for its wood and nuts. It is the third broad-leaved tree species in France in forest area (750,000 ha) and in 2016, accounted for 5% of land used for fruit production (FranceAgriMer 2017). With an annual production of 7,000-9,000 tons in the last 10 years, France is the fifth European producer (FAO 2018). Sweet chestnut has been intensively cultivated in coppices and orchards for centuries in France. However, since the beginning of the 18^th^ century, it has suffered from abandonment, leading to a sharp decrease in production (Pitte 1986; Sauvezon et al. 2000). Many landraces and associated knowledge were lost. In the early 20^th^-century, the Laffite tree nursery in Basque Country started to breed hybrid chestnut followed in the 1960s by the French National Institute for Agricultural Research (INRA) (Pereira-Lorenzo et al., 2016). These breeding programs aimed at developing interspecific hybrids resistant to ink disease caused by a *Phytophtora* fungus, by crossing two Asian tolerant species, *Castanea crenata* and *Castanea mollissima*, with local landraces from regions with an oceanic climate. These hybrids are now mainly used for fruit production and as rootstock (particularly Marigoule and Bouche de Bétizac varieties, and more recently BelleFer). However, they are not adapted to continental and Mediterranean conditions (Martin et al. 2017; Míguez-Soto et al. 2019). Their fruit quality has also been criticized by some growers and by chestnut lovers, particularly in comparison with landraces. Action was thus taken by these actors, involving survey of old chestnut trees, phenotypic observations and the establishment of conservatory orchards.

Strong geographical structure was reported in wild populations in Italy, Spain, Greece and Turkey (Mattioni et al. 2013). A study of wild chestnut in Spain, Italy and Greece (Fernández-Cruz and Fernández-López 2016) found two main gene pools in Europe, and another study of wild, natural or naturalized populations across Europe (Mattioni et al. 2017), found three. These findings agree with evidence of spontaneous establishment originating from the Last Glacial Maximum refugia in the North of the Iberian, Italian and Balkan peninsulas, and in northern Anatolia (Krebs et al. 2004, 2019; Roces-Díaz et al. 2018). In southern France, there is possible evidence for chestnut refugia in palaeo-botanical data (Krebs et al. 2019). The preferred hypothesis is therefore that most pre-cultivation Castanea in France are the result of the spontaneous spread of the species from neighboring southern European refugia, i.e. in Spain and Italy. However, the most recent genetic analyses conducted exclusively on French populations were published in the 1990s on wild chestnut and at a regional scale (Frascaria et al. 1991, 1992; Frascaria and Lefranc 1992) the and the results obtained in the CASCADE project (Eriksson et al. 2005) have not yet been published (T. Barreneche pers. com.). Mattioni et al. (2008) compared naturalized, coppice and orchard populations in Italy, Greece, Spain, the UK and France, and showed differences in within-population genetic parameters between fruit orchards and other types of chestnut management. This result implies that long-term management techniques can influence the genetic makeup of the populations. Differences between and within countries have also been reported (Pereira-Lorenzo et al. 2016). For these reasons, specifically French, finer-scale sampling of both wild (forest) and cultivated chestnut trees (orchards and alignments) was needed to help distinguish between natural and anthropogenic evolutionary factors.

In terms of sampling, many authors have genotyped tree collections *ex situ*, i.e., in conservatories (Martín et al. 2010a), and *in situ* (Pereira-Lorenzo and Fernandez-Lopez 1997; Gobbin et al. 2007; Martin et al. 2010b; Pereira-Lorenzo et al. 2010, 2019; Beccaro et al. 2012; Mellano et al. 2012, 2018; Beghè et al. 2013; Quintana et al. 2014; Fernández-López and Fernández-Cruz 2015). In this study, we used both in- and ex-situ sources to assess the known and currently used genetic diversity of sweet chestnut. As a result, we often sampled several individuals belonging to the same landrace. Hereafter, we use the term “landrace” as defined by Villa et al. (2005) rather than “variety” or “cultivar”, as it better covers the variety of sampling situations we encountered in the field. However, we do use the term “cultivar” when known cultivars were encountered.

The main aims of this work were to assess (i) the genetic diversity of wild and cultivated chestnut in most of its range in France, (ii) their genetic structure, notably in relation with the sampling regions, and (iii) relations between French chestnut and its neighbors in Spain and Italy. For this purpose, we sampled natural chestnut populations, ancient grafted chestnut identified *in situ* by local partners and *ex situ* local landraces in conservatories in the main nut-producing regions and in most of the distribution of natural chestnut forests in France. We used microsatellite markers from the EU chestnut database to genotype all sampled trees at 13 SSRs and a subset at 24 SSRs (Pereira-Lorenzo et al. 2017). By also including Iberian samples cited in the Pereira-Lorenzo et al. publication, we also provide some evidence for the origin of the trees we sampled.

Fixation of genotypes by grafting from spontaneous chestnut, or “instant domestication” as defined by (Harris et al. 2002), is reported in the literature (Aumeeruddy-Thomas et al. 2012), and was recently documented in Italy and Spain (Pereira-Lorenzo et al. 2019). As a working hypothesis, this suggested a possible lack of genetic structure between wild and cultivated chestnut. It is common knowledge that grafts and nuts travel by means of markets, historically via occupational travelers such as glass blowers (Pitte 1986) and now via local and internet-mediated exchange fairs. However, the extent and impact of this phenomenon on the genetic structure of cultivated chestnut was previously unknown in France. We hypothesized that it is sufficiently frequent to have a significant impact, leading to a low genetic structure of cultivated chestnut in France. As reported in (Pereira-Lorenzo et al. 2019), we also expected to find a high overall genetic diversity, but without marked differences between the wild and cultivated sets. In addition, we expected *in situ* local landraces to be multiclonal due to repeated grafting over the centuries and the accumulation of mutations or the use of seedlings from the landrace.

## MATERIALS AND METHODS

Terminology: we avoid the use of “population” and instead use “sampling region” to describe a geographically or socially meaningful region where a non-profit association has prospected and conserved chestnut, or a group of sampling sites located close by. We use “genetic cluster” to denote a cluster of genotyped trees resulting from the analysis of their genetic structure. “Chestnut type” is used as a category with two levels, “forest” and “cultivated”.

### 2.1. Sampling scheme

To characterize and understand the genetic diversity and population structure of the European chestnut (*Castanea sativa* Mill.) in France, we genotyped 693 trees in 16 sampling regions in both forest and cultivated areas. Table 1 lists sampling details and Figure 1 shows the location of the sampling regions (GPS of sampled trees are available upon request).

**Table 1:**
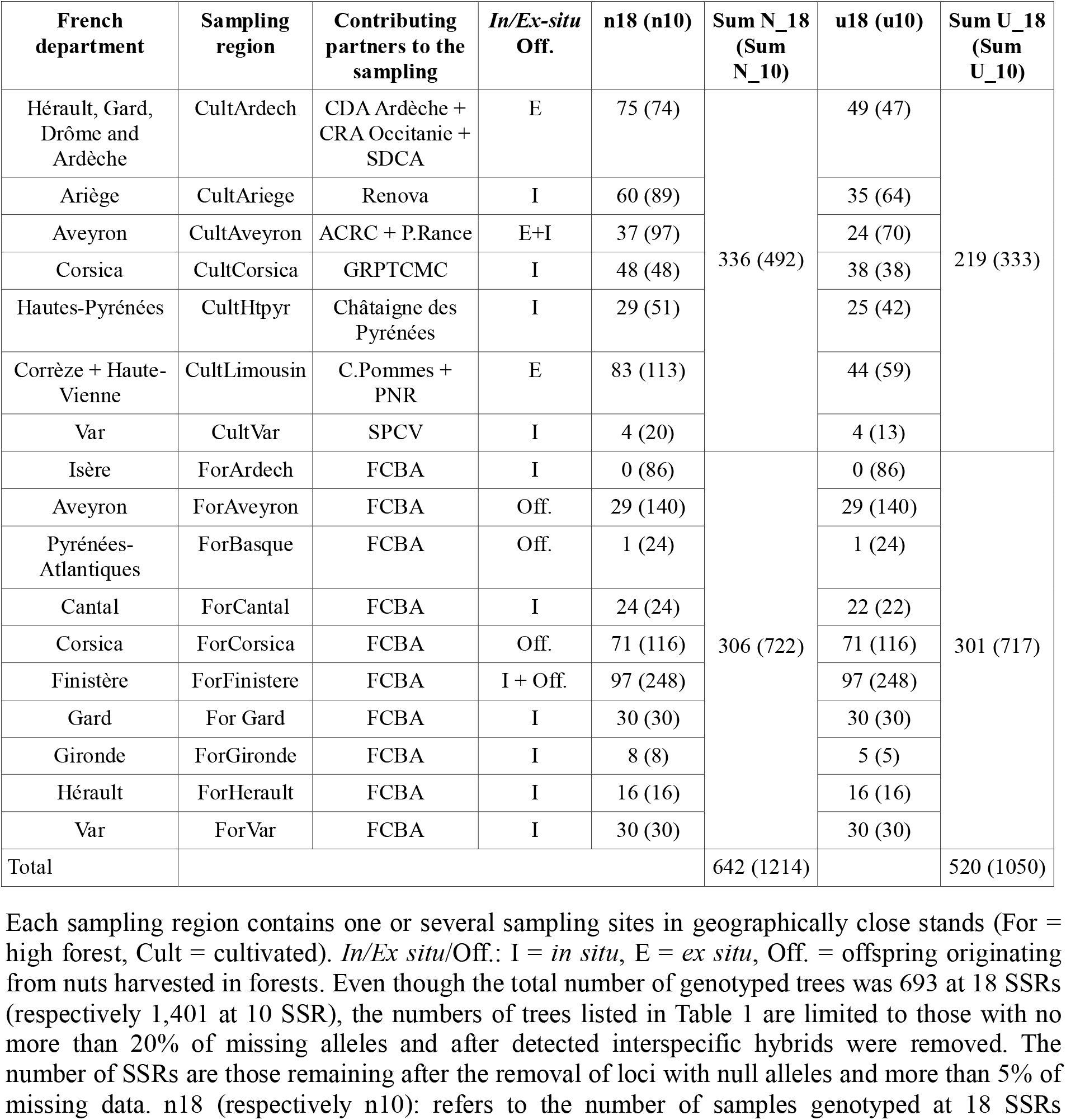
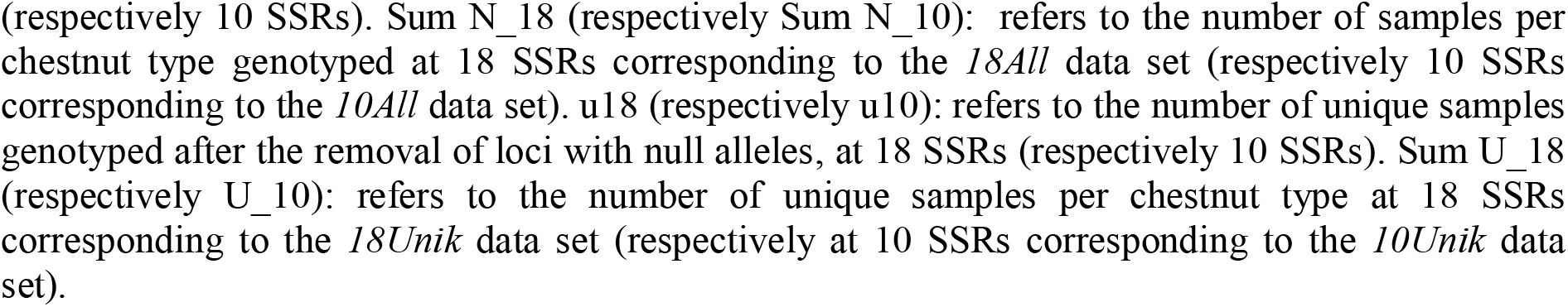
Description of the sampling regions and sampled trees

| French department | Sampling region | Contributing partners to the sampling | In/Ex-situ Off. | n18 (n10) | Sum N_18 (Sum N_10) | u18 (u10) | Sum U_18 (Sum U_10) |
| --- | --- | --- | --- | --- | --- | --- | --- |
| Hérault, Gard, Drôme and Ardèche | CultArdech | CDA Ardèche + CRA Occitanie + SDCA | E | 75 (74) | 336 (492) | 49 (47) | 219 (333) |
| Ariège | CultAriege | Renova | I | 60 (89) |  | 35 (64) |  |
| Aveyron | CultAveyron | ACRC + P.Rance | E+I | 37 (97) |  | 24 (70) |  |
| Corsica | CultCorsica | GRPTCMC | I | 48 (48) |  | 38 (38) |  |
| Hautes-Pyrénées | CultHtpyr | Châtaigne des Pyrénées | I | 29 (51) |  | 25 (42) |  |
| Corrèze + Haute-Vienne | CultLimousin | C.Pommes + PNR | E | 83 (113) |  | 44 (59) |  |
| Var | CultVar | SPCV | I | 4 (20) |  | 4 (13) |  |
| Isère | ForArdech | FCBA | I | 0 (86) | 306 (722) | 0 (86) | 301 (717) |
| Aveyron | ForAveyron | FCBA | Off. | 29 (140) |  | 29 (140) |  |
| Pyrénées-Atlantiques | ForBasque | FCBA | Off. | 1 (24) |  | 1 (24) |  |
| Cantal | ForCantal | FCBA | I | 24 (24) |  | 22 (22) |  |
| Corsica | ForCorsica | FCBA | Off. | 71 (116) |  | 71 (116) |  |
| Finistère | ForFinistere | FCBA | I + Off. | 97 (248) |  | 97 (248) |  |
| Gard | For Gard | FCBA | I | 30 (30) |  | 30 (30) |  |
| Gironde | ForGironde | FCBA | I | 8 (8) |  | 5 (5) |  |
| Hérault | ForHerault | FCBA | I | 16 (16) |  | 16 (16) |  |
| Var | ForVar | FCBA | I | 30 (30) |  | 30 (30) |  |
| Total |  |  |  |  | 642 (1214) |  | 520 (1050) |

Each sampling region contains one or several sampling sites in geographically close stands (For = high forest, Cult = cultivated). *In/Ex situ*/Off.: I = *in situ*, E = *ex situ*, Off. = offspring originating from nuts harvested in forests. Even though the total number of genotyped trees was 693 at 18 SSRs (respectively 1,401 at 10 SSR), the numbers of trees listed in Table 1 are limited to those with no more than 20% of missing alleles and after detected interspecific hybrids were removed. The number of SSRs are those remaining after the removal of loci with null alleles and more than 5% of missing data. n18 (respectively n10): refers to the number of samples genotyped at 18 SSRs
(respectively 10 SSRs). Sum N<sub>18</sub> (respectively Sum N<sub>10</sub>): refers to the number of samples per chestnut type genotyped at 18 SSRs corresponding to the *18All* data set (respectively 10 SSRs corresponding to the *10All* data set). u<sub>18</sub> (respectively u<sub>10</sub>): refers to the number of unique samples genotyped after the removal of loci with null alleles, at 18 SSRs (respectively 10 SSRs). Sum U<sub>18</sub> (respectively U<sub>10</sub>): refers to the number of unique samples per chestnut type at 18 SSRs corresponding to the *18Unik* data set (respectively at 10 SSRs corresponding to the *10Unik* data set).

**Fig. 1:**
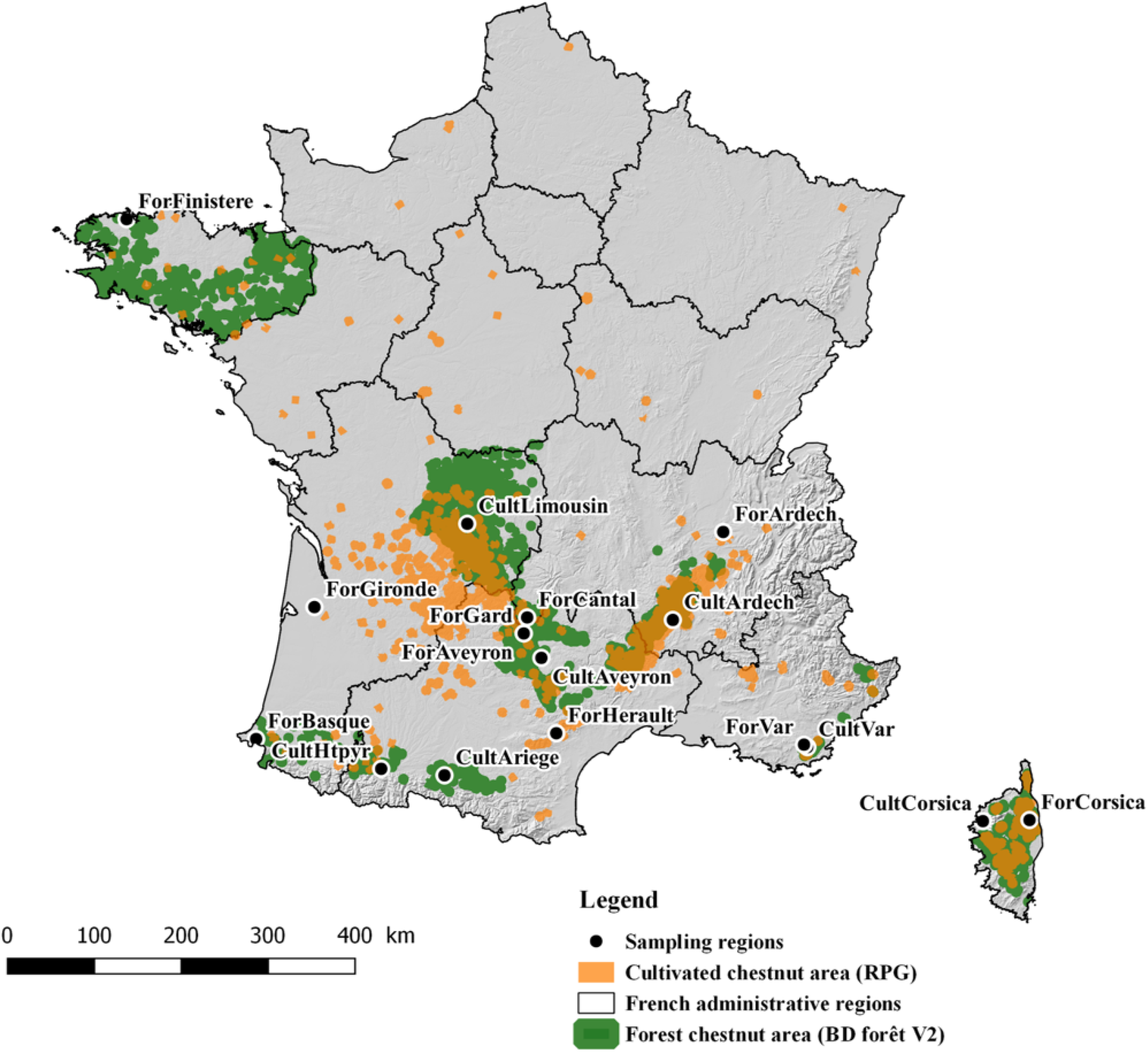
Map of sampling regions. The distribution of chestnut forest areas where chestnut accounts for at least 75% of the leaf cover, which represents about 50% of the total chestnut-comprising forest area in France (IGN 2007) (in green, and the distribution of cultivated chestnut and orchards (IGN 2016) (in orange). Each dot represents a sampling region where chestnut forest or orchard is present (this qualitative information does not reflect the relative areas or number of trees).

### 2.2. Geographical sampling

In forest stands, trees were chosen randomly, located several dozen meters apart in the middle of forest patches. Their exact locations were recorded by GPS. In Brittany, Auvergne-Rhône-Alpes, Occitanie, Provence-Alpes-Côtes d’Azur (PACA) and Corsica, mature leaves were sampled and immediately enclosed in plastic bags with silicagel. In Gironde, dormant buds were sampled from trees close to the laboratory to facilitate frequent re-sampling when assessing the accuracy of genotyping protocols. In Corsica, nuts and dried leaves were also sampled in the field, whereas cultivated chestnut was provided as DNA extract. Whenever we sampled offspring as groups of half sib fruits, we also sampled leaves from their mothers. Nuts harvested in the Finistère, Corsica, Basque Country and Aveyron forest sampling regions were germinated and sown in the greenhouse.

### 2.3. Expert-based sampling

Field surveys of cultivated chestnut were conducted in 2016-2017 in collaboration with producer and amateur organizations. In 2016, we focused our sampling effort on the landraces they knew and were interested in. In 2017, we expanded sampling to most known landraces and grafted trees, supplemented by random sampling in a few chestnut orchards. Associative conservatories were also sampled. We sampled several chestnut trees that had the same name to test the genetic diversity of landraces. When attributing sampled trees to a given landrace, when known, we followed the field expert’s determination.

### 2.4. SSR genotyping

A total of 693 trees were genotyped at 24 SSRs and 1401 trees at 13 SSRs. Total genomic DNA was extracted from fresh leaves, silica-dried leaves or dormant buds using the DNeasy 96 Plant kit (Qiagen, Hilden Allemagne). Twenty-four SSR markers previously selected to study chestnut genetic diversity were used for this study (Buck et al. 2003; Gobbin et al. 2007; Kampfer et al. 1998; Marinoni et al. 2003; Steinkellner et al. 1997) based on the protocol of Pereira-Lorenzo et al. (2017). We amplified these 24 SSRs into 5 multiplex and 2 singleplex PCRs using one of the FAM, NED, PET, VIC fluorophore-labeled primers (PE Applied Biosystems, Warrington, UK) modified following (Pereira-Lorenzo et al. 2017, 2019). The PCR final reaction volume was 15 μl (7.5 μl of QIAGEN Multiplex Master Mix, 0.075 to 0.3 μM of each primer, 4 to 4.9 μl RNase Free Water and 2 μl of ADN at 5-10 ng/μl). The amplification conditions were 95°C for 5 min, followed by 30 cycles at 95°C for 30 s, annealing at a specific temperature depending on the multiplex set, for 1.5 min, and 1 min at 72°C, and final extension at 60 °C for 30 min. Negative controls were included in all PCR reactions to enable detection of cross contamination of the samples. Amplifications at 13 SSRs corresponded to sets 1, 2 and 3. Amplifications at 24 SSRs corresponded to all sets. The descriptions of all sets are displayed in Table 2. Amplification products were diluted with water, 2 μl of the diluted amplification product was added to 0.12 μl of 600LIZ size standard (Applied Biosystems, Foster City, USA) and 9.88 μl of formamide.

**Table 2:**
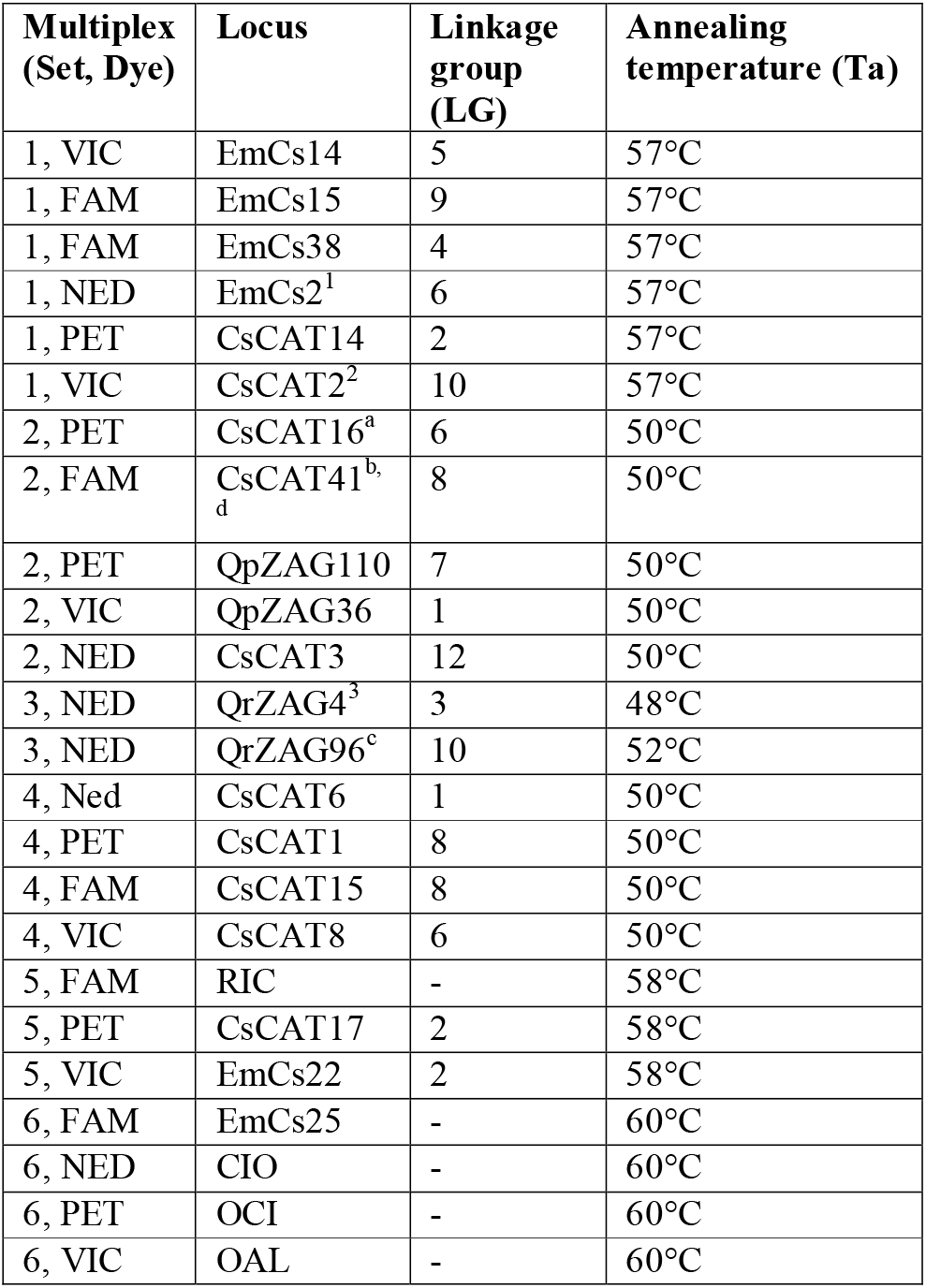
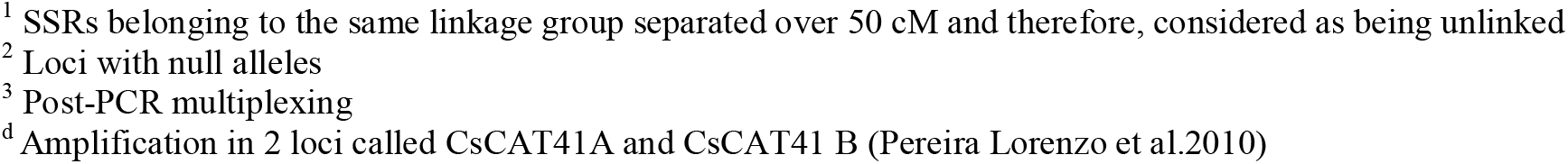
SSRs markers and multiplex.

| Multiplex (Set, Dye) | Locus | Linkage group (LG) | Annealing temperature (Ta) |
| --- | --- | --- | --- |
| 1, VIC | EmCs14 | 5 | 57°C |
| 1, FAM | EmCs15 | 9 | 57°C |
| 1, FAM | EmCs38 | 4 | 57°C |
| 1, NED | EmCs2<sup>1</sup> | 6 | 57°C |
| 1, PET | CsCAT14 | 2 | 57°C |
| 1, VIC | CsCAT2<sup>2</sup> | 10 | 57°C |
| 2, PET | CsCAT16<sup>a</sup> | 6 | 50°C |
| 2, FAM | CsCAT41<sup>b, d</sup> | 8 | 50°C |
| 2, PET | QpZAG110 | 7 | 50°C |
| 2, VIC | QpZAG36 | 1 | 50°C |
| 2, NED | CsCAT3 | 12 | 50°C |
| 3, NED | QrZAG4<sup>3</sup> | 3 | 48°C |
| 3, NED | QrZAG96<sup>c</sup> | 10 | 52°C |
| 4, Ned | CsCAT6 | 1 | 50°C |
| 4, PET | CsCAT1 | 8 | 50°C |
| 4, FAM | CsCAT15 | 8 | 50°C |
| 4, VIC | CsCAT8 | 6 | 50°C |
| 5, FAM | RIC | - | 58°C |
| 5, PET | CsCAT17 | 2 | 58°C |
| 5, VIC | EmCs22 | 2 | 58°C |
| 6, FAM | EmCs25 | - | 60°C |
| 6, NED | CIO | - | 60°C |
| 6, PET | OCI | - | 60°C |
| 6, VIC | OAL | - | 60°C |

Genotyping was performed partly on an ABI 310 capillary sequencer (Applied Biosystems, Foster City, CA, USA) at the Xylobiotech FCBA facility of Cestas-Pierroton with further work on an ABI 3500 XL capillary sequencer (Applied Biosystems, Foster City, CA, USA) at the CIRAD GenSeqUM platform in Montpellier, France. Allele sizes were read independently by two investigators using GENEMAPPER 4.1 and 5.0 respectively (Applied Biosystem, Foster City, USA). The output files in the fsa format were made compatible for GENEMAPPER 4.1 using a Python script from the Montpellier platform.

### 2.5. DATA ANALYSIS

#### 2.5.1. Data filtering: Detection of interspecific hybrids, clonal groups and null alleles

All individuals with more than 20% of missing alleles were removed along with individuals showing Asiatic alleles. Asiatic genotypes present initially in our data and Asiatic alleles identified in a previous study (Pereira-Lorenzo et al. 2010) were used to detect exotic germplasm in the European populations.

The presence of uninformative loci was tested with the informloci function in the R/poppr package version 2.8.3 (Kamvar et al. 2015; Kamvar et al. 2014) in both data sets. The percentages of missing data were obtained using the info_table function in R/poppr. The frequency of null alleles per locus was calculated with the R/PopGenReport package version 3.0.4 (Adamack and Gruber 2014) based on Brookfield formula (Brookfield 1996). Following (Lassois et al. 2016), we discarded loci with more than 10% of null alleles. After removing loci, the genotype curve function implemented in the R/poppr. was applied to both data sets to determine the minimum number of loci necessary to discriminate between individuals. Redundant genotypes were searched within each sampling region to identify multi-locus genotypes (MLGs) for each data set, using the clonecorrect function in R/poppr. When detected these redundant MLGs were removed. Note that some MLGs may be redundant between sampling regions but those were kept for the rest of the analyses as they provide information on population structure.

#### 2.5.2. Genetic diversity

The observed number of alleles (Na) and observed heterozygosity (Ho) were calculated at each locus across sampling regions using the summary function in the R/adegenet package 2.1.1 (Jombart 2008). The effective number of alleles (Ne) was calculated using the expected heterozygosity (He) from the summary function for genind object in R/adegenet, with Ne=1/(1-He). The Fst and corrected Fst (Fstp), Fis and Dest per locus (Jost 2008; Nei 1987) were calculated using the basic.stats function in the R/hierfstat package version 0.04-22 (Goudet 2005). The poppr function in R/poppr was used to report other basic statistics per sampling region including the Shannon-Weiner diversity index (H), the index of association (Ia), and the standardized index of association (rbarD) (Agapow and Burt 2001). The significance of Ia and rbarD were tested with 1000 permutations, shuffling the genotypes at each locus while maintaining the heterozygosity and allelic structures. Deviation from the Hardy-Weinberg equilibrium (HWE) was tested on both loci and populations with 1000 permutations using the hw.test function in the R/pegas package version 0.11 (Paradis 2010). The Chi2 statistic was calculated over the entire data set and two p values were computed, one analytical and one derived from 1000 Monte-Carlo permutations.

#### 2.5.3. Population structure

For each data set, three methods were compared to find genetic clusters, by varying the number of clusters, K, between 1 and 15: STRUCTURE (Pritchard et al, 2000; Falush et al, 2003), discriminant analysis of principal components (DAPC: Jombard et al, 2010) and sparse nonnegative matrix factorization (SNMF: Frichot et al, 2014). For the first method, the STRUCTURE software v2.3.4 was launched thirty times for each K value using the admixture model with unlinked loci and correlated frequencies (other parameters: usepopinfo=0, popflag=0, burn-in=200,000, MCMC iterations=200,000). The number of clusters was determined using the DeltaK metric from structureHarvester v0.6.94 (Evanno et al, 2005). At the best K, the assignment probabilities of each individual to each cluster (Q matrix) were averaged over the 30 runs using a suitable permutation to handle label switching. The permutations were obtained with the CLUMPP software v1.1.2 using the LargeKGreedy algorithm with 2000 repeats (Jakobsson and Rosenberg, 2007). For the second method, SSR genotypes were first transformed by a principal component analysis (PCA), followed by the K-means algorithm applied to the principal components using the find.clusters function in R/adegenet v2.1. The number of clusters was determined using the BIC. The DAPC was then performed at the best K with the number of principal components to keep chosen by cross-validation using the xvalDapc function in R/adegenet with 30 repetitions and a maximum of 80 PCs. For the third method, the snmf function of R/LEA v2.0 was launched with 5 repetitions. The number of clusters was determined using the cross-entropy criterion. For all methods, the clusteredness metric was computed to assess the extent to which a randomly chosen individual is inferred to have ancestry in only one cluster (Rosenberg et al, 2005). The threshold above which individuals were considered as “strongly assigned” was set to 0.8 (qI ≥ 80%).

Hierarchical analysis of molecular variance (AMOVA, Excoffier and Smouse 1992) as implemented in the poppr.amova function in R/poppr was performed using all loci with less than 5% missing data on the preset hierarchy of chestnut types and sampling regions, and on genetic clusters. Fis, pairwise Fst and hierarchical F-statistics were calculated, and 95% confidence intervals were obtained by bootstrapping with 1000 samples over loci using the boot.ppfis, boot.ppfst and boot.vc functions. Differences between hierarchy levels were tested by randomization with the function randtest in the R/ade*4* package version 1.7-13 (Excoffier and Smouse 1992; Chessel et al. 2004). Some components of covariance could have slightly negative estimates due to the absence of significant genetic structure at the corresponding hierarchical level (FAQ List for Arlequin 2.000).

#### 2.5.4. Reproducibility

To facilitate method reproducibility (Goodman et al. 2016), most of our analyses were performed in R (R Core Team 2019); the scripts are available at https://data.inra.fr/privateurl.xhtml?token=8c03a83c-be4d-4984-972f-7808558b4539.

## 3. RESULTS

### 3.1. Data filtering: Detection of interspecific hybrids, clonal groups and null alleles

For the 24 SSRs data set, 13 genotyped trees with more than 20% of missing alleles were filtered out (respectively 32 trees at 13 SSRs). Three trees genotyped twice were filtered out at 24 SSRs (respectively 18 trees at 13 SSRs). Over all our data sets, Asiatic alleles were scored as following: CsCAT41A-196, EmCs15-76, EmCs2-152, QpZAG36-205, QrZAG4-118, QrZAG96-154, EmCs14-135, CsCAT14-136, CsCAT6-138, CsCAT1-188, CsCAT17-135, CIO-160, OCI-160, OCI-167, OCI-169 and OAL-307. For the 24 SSRs data set, 26 interspecific trees were filtered out, 11 were known beforehand and 15 were detected out of known Asiatic alleles (respectively 127 trees at 13 SSRs : 18 were known beforehand and 119 were detected out of known Asiatic alleles). Most of the detected interspecific trees were from ForBasque. Nine Spanish trees were filtered out at 24 SSRs (respectively 10 trees at 13 SSRs). A total of 51 trees were discarded at 24 SSRs (respectively 187 trees at 13 SSR). For the 24 SSRs data set, 642 trees genotyped (respectively 1,214 at 13 SSRs) remained for further analysis (Table 1).

CsCAT41 is known to amplify two sites: CsCAT41A and CsCAT41B. CsCAT41A was less polymorphic, with only 2 alleles detected in our analyses out of 3 already known (Pereira-Lorenzo et al, 2010): CsCAT41A-196 which matches with CsCAT41A-200 in Pereira-Lorenzo et al (2010) and CsCAT41A-199 which matches with CsCAT41A-202 in Pereira-Lorenzo et al (2010). CsCAT41A-188 in Pereira-Lorenzo et al (2010) did not came out in our data set. CsCAT41A was removed from the subsequent analyzes. CsCAT41B was more polymorphic, with 11 alleles (210-233) in the data set with 24 SSRs and 13 alleles (210-237) in the data set with 13 SSRs.

After filtering for null alleles, EmCs38, CIO and EmCs25 were discarded from the data set at 24 SSRs (respectively EmCs38 at 13 SSRs) (Online Resource 1). EmCs14, EmCs2 and EmCs25 had more than 5% of missing alleles and were discarded from the data set at 24 SSRs (respectively QrZAG4 at 13 SSRs). The non-phylogenetically informative loci EmCs14 was discarded from both data sets (minor allele frequency MAF <0.01).

The resulting data sets had 18 SSRs, hereafter called *18All*, and 10 SSRs, hereafter called *10All*. Redundant multi-locus genotypes (MLGs) were then detected in each sampling region, as they *could be the result of both practices (grafting) and sampling choices They had to be removed to avoid the artefactual detection of genetic structure resulting from the sampling strategy*. The resulting data sets (Table 1) are called *10Unik* (1050 trees) and *18Unik* (520 trees). 4 MLGs among sampling regions were detected at 24 SSRs (respectively 9 at 13 SSRs) but were not discarded as they provide information on populations structure. In both data sets, the discriminating power of the polymorphic markers to differentiate between genotypes was sufficient to discriminate all individuals irrespective of the number of loci and individuals (Online Resource 2).

### 3.2. Description of SSR diversity per sampling region

The 18 SSRs analyzed in this study varied greatly in allele diversity (Online Resource 3). The *18Unik* data set (respectively *10Unik*) had a total of 179 alleles (respectively 112) with an average of 9.9 alleles per locus (respectively 11.2). This ranged from 2 for QrZAG4 to 31 for CsCAT3 (respectively 3 for EMCs2 to 33 for CsCAT3). In terms of expected heterozygosity (He), QrZAG4 showed the lowest diversity with 0.17 in *18Unik* (respectively EMCs2 with 0.66 in *10Unik*) and CsCAT6 the highest diversity with 0.86 in *18Unik* (respectively CsCAT3 with 0.85 in *10Unik*). The within-population inbreeding coefficient (Fis) ranged from −0.427 to 0.153 in *18Unik* (respectively −0.431 to 0.169 in *10Unik*), with a mean of −0.102 in *18Unik* (respectively −0.031 in *10Unik*). In *18Unik*, only EmCs15, CsCAT16, QpZAG110, QrZAG4, CsCAT1, CsCAT15, RIC and OCI were in the HWE. Across all sampling regions, in *10Unik,* it was not possible to reject the HWE for CsCAT3 and QpZAG110 (Online Resource 4). When tested per sampling region, only ForGard and ForBasque were in the HWE in both data sets. ForAveyron was in the HWE only in *10Unik*. Moreover, in both data sets, HWE was rejected for all SSR loci in at least one sampling region except OCI in *18Unik*

### 3.3. Redundant diversity among sampling regions and no differentiation between chestnut types

Genetic diversity indices calculated for each sampling region genotyped at 18 SSRs without MLGs are listed in Table 3 (results at 18 SSRs are presented in Online Resource 5). Note that the aim of sampling ForGironde was not to be representative of the region, but to facilitate resampling. Moreover, in the *18Unik* data set ForBasque had a single individual. Therefore, diversity and differentiation are discussed excluding ForGironde and ForBasque in the *18Unik* data set, and excluding ForGironde in the *10Unik* data set. The highest effective number of alleles per sampling region was found in the Finistère forest sampling regions in *18Unik* (ForFinistere, northwestern France) and the lowest was found in the cultivated sampling region in Var (CultVar, south east of France). The mean observed heterozygosity was 0.671 and the mean expected heterozygosity was 0.605. The sampling regions with the lowest (respectively highest) observed heterozygosity were ForAveyron, the forest sampling region in Aveyron (respectively CultVar). The sampling regions with the lowest (respectively highest) expected heterozygosity were CultVar (respectively the forest sampling regions in Finistère, ForFinistere). Excluding ForGironde, positive and significant inbreeding (Fis) were found in the cultivated sampling region in Ariège (CultAriege) and in the forest sampling region in Ardèche (ForArdech). The highest Ia and rbarD were found in CultVar and the lowest were found in ForFinistere. The results of AMOVA (Table 4 and Online Resource 6), revealed no substantial difference in structure in chestnut type between forest stands and cultivated orchards: the variance component did not significantly differ from zero. Moreover, althoughthe Fct had a confidence intervals excluding zero, it is very close to zero. Instead, more than 80% of the variance was found within each sampling region. At a threshold of 0.001, we rejected the null hypothesis of panmixia, both among sampling regions within chestnut types and within sampling regions. Among sampling regions within chestnut types, the Phi test statistic of the AMOVA indicated greater variance than expected under the null hypothesis. This suggested an underlying structure at this hierarchical level that was confirmed by a positive bootstrap-derived confidence interval for Fst (0.088-0.11) Within sampling regions, the Phi test statistic indicated lower variance than expected under the null hypothesis. This suggested some inbreeding at this hierarchical level confirmed by a negative bootstrap-derived confidence interval for Fis, although very close to zero. Note that in the *10Unik* data set, a bootstrap-derived confidence interval for Fis included zero (Online Resource 6).

**Table 3:**
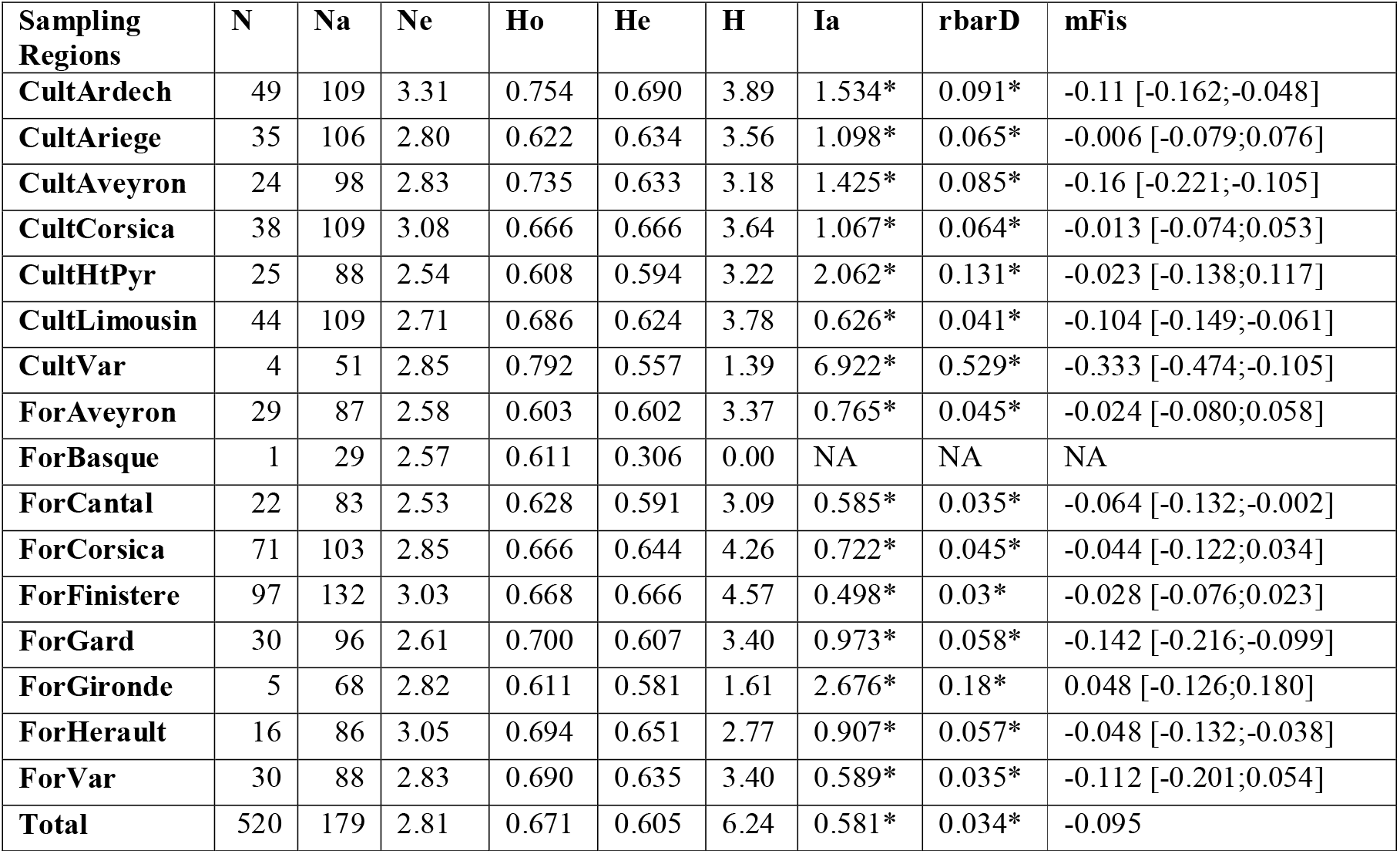
Genetic diversity indices for 16 French sampling regions at 18 loci without MLGs (*18Unik* data set) N: number of unique individuals genotyped per sampling region; Na: number of alleles; Ne: mean number of effective alleles; Ho: observed heterozygosity; He: expected heterozygosity; H: Shannon-Weiner diversity index; Ia: index of association; rbarD: standardized index of association; Fis: inbreeding coefficient, with 95% confidence interval (CI). Asterisks indicate significant *p* values at the 0.001 threshold. The “Total” row contains the sum of N, total Na and total H, and the mean for the other indices.

**Table 4:** Hierarchical AMOVA and F-statistics for 16 French sampling regions at 18 loci without MLGs (*18Unik* data set). df: degree of freedom, Alter: alternative hypothesis, 95% confidence intervals, ***: p value ≤ 0.001

| Source of variation | df | Variance component | % variation | *p* value | Alter. | F statistic |
| --- | --- | --- | --- | --- | --- | --- |
| Among chestnut type | 1 | -0.169 | -1.54 | 0.886 | greater | Fct -0.01 [-0.013; -0.005] |
| Among sampling regions within chestnut types | 14 | 1.740 | 15.81 | 0.001*** | greater | Fst 0.099 [0.088; 0.11] |
| Within sampling regions | 504 | 9.435 | 85.73 | 0.001*** | less | Fis -0.036 [-0.062; -0.005] |
| Total | 509 | 11.006 | 100.00 |  |  | Fit 0.058 [0.03; 0.093] |

### 3.4. Highly admixed genetic structure

In addition to analyzing genetic diversity per sampling region, we also evaluated the overall genetic structure to detect genetic clusters, if any, and to assess their congruence with respect to the sampling regions.

The number of genetic clusters was determined using three methods, each with its own criterion: DeltaK for STRUCTURE, BIC for DAPC and cross-entropy for SNMF. For all data sets, BIC and cross-enthropy started by decreasing sharply (Online Resource 7), demonstrating the presence of genetic structure. However, the signal was not so clear, making the choice of the number of genetic clusters rather difficult. By contrast, the signal of DeltaK was clear for all data sets. Therefore, we chose to display in the main and supplementary material the results of STRUCTURE rather than the other methods.

On the *18Unik* data set (respectively *10Unik*), the most likely number of clusters, according the DeltaK criterion (Online Resource 7), gave the highest value for K=2 (respectively K=6). We chose to display the results at K=2 and K=6 for the *10Unik* and *10Unik with Spanish samples* data sets in order to allow comparison with the *18Unik and 18Unik with Spanish samples* data set.

Reconstructed panmictic population (RPP) were numbered according the clustering of the *18Unik* data set at K=2 (RPP1 and RPP2) and the clustering of the *10Unik* data set was named in reference to the K=2 clustering (RPP1a, RPP1b, RPP2a, RPP2b, RPP2c and RPP2d). In tables and online material, the numbers of clusters were kept together with the RPPs in order to help the reader to follow the results from both the manuscript and the files hosted on INRAEserver.

On the *18Unik* (respectively *10Unik*) data set, for K=2, 85% (respectively 85%) of the individuals from Corsica, Var and Ardèche were grouped in reconstructed panmictic population 1 (RPP1) and 68% (respectively 62%) of individuals from all sampling regions except Var (respectively except the forest region in Var) were grouped in RPP2, pointing to an overall admixed genetic structure in our sample. On the *10Unik* data set, for K=6 95% of the individuals from Var and 20% of individuals from Ardèche were in RPP1a, 94% of the individuals from Corsica were grouped in RPP1b, 70% of individuals from Limousin and Finistère (west of France) were grouped in RPP2a, 71% of individuals from Aveyron and Cantal (south-west of Massif Central) were grouped in RPP2b, 68% of individuals from Ardèche were grouped in RPP2c, 80% of individual from Ariège, Hautes-Pyrénées and Basque Country (French Pyrennées) and 69% of individuals from Hérault were grouped in RPP2d.

The overall admixed genetic structure in our sample was confirmed by the relatively low pairwise Fst calculated between clusters, as can be seen in Figure 2 and Online Resource 8: 0.09 between RPP1 and RPP2 when ql≥80% on the *18Unik* (respectively 0.088 on the *10Unik*) data set. When K=6 and ql≥80% on the *10Unik* data set, the highest Fst values were found between Var and Ardèche (RPP1a) and Aveyron (RPP2b), West of France (RPP2a) and many sampling regions (RPP2d). The lowest Fst values were found between RPP1a, RPP2b and RPP2d. Taking into account the confidence interval, the genetic differentiation between Var and Ardèche (RPP1a) and Corsica (RPP2b) is of the same magnitude than the genetic differentiation between Var and Ardèche (RPP1a) and Aveyron (RPP2b), West of France (RPP2a) and many sampling regions (RPP2d).On the *18Unik* (respectively *10Unik*) data set, eighty-five individuals out of 520, i.e., 16% (respectively 564 out of 1,050, i.e., 54%) had a ql < 80%. A hierarchical AMOVA performed on the strongly-assigned individuals (ql ≥80%) of the *18Unik* data set (respectively *10Unik*) corroborated this finding (Table 5 and Online Resource 6):86.6% of the variance (respectively 81.9%) was found among samples within clusters. For the *18Unik* (respectively *10Unik*) data set, the variance component between clusters was 13.4% (respectively 18.1%) and the Fst was of 0.09 (respectively 0.125), indicating a relatively low but significant genetic structure (Table 5 and Online Resource 6). When the inbreeding coefficient was calculated per cluster on strongly-assigned individuals (Table 6 and Online Resource 9), the 95% confidence interval of both clusters included zero. The mean observed heterozygosity was 0.673 for *18Unik* (respectively 0.657 for *10Unik*) and the mean expected heterozygosity was 0.681 (respectively. 0.654 for *10Unik*).

**Table 5:**
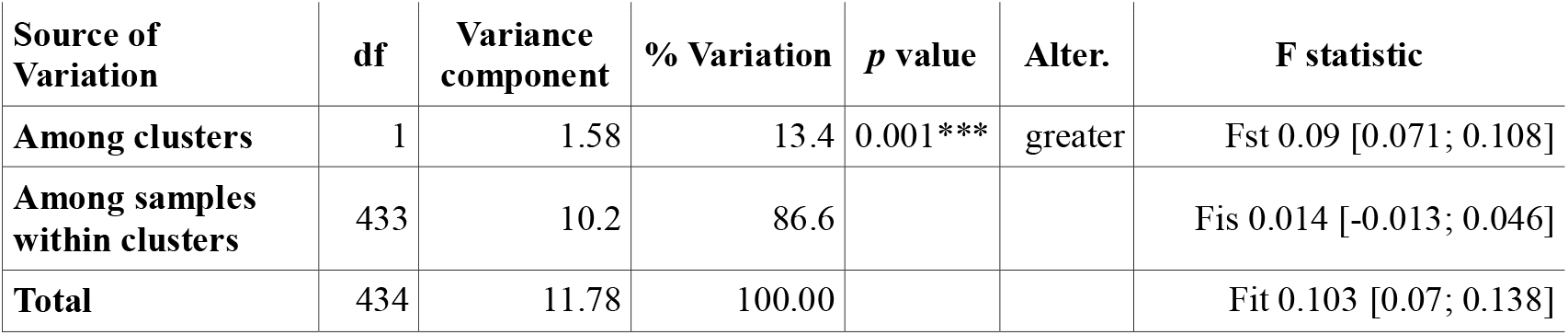
Hierarchical AMOVA and F-statistics for two genetic clusters at 18 loci without MLGs, on strongly assigned individuals (ql ≥ 80%) df: degrees of freedom, Alter: alternative hypothesis, 95% confidence intervals, ***: p value ≤ 0.001

| Source of Variation | df | Variance component | % Variation | *p* value | Alter. | F statistic |
| --- | --- | --- | --- | --- | --- | --- |
| Among clusters | 1 | 1.58 | 13.4 | 0.001*** | greater | Fst 0.09 [0.071; 0.108] |
| Among samples within clusters | 433 | 10.2 | 86.6 |  |  | Fis 0.014 [-0.013; 0.046] |
| Total | 434 | 11.78 | 100.00 |  |  | Fit 0.103 [0.07; 0.138] |

**Table 6:** Within-cluster genetic variability at 18 loci without MLGs, on strongly assigned individuals (ql ≥ 80%) N: number of unique individuals genotyped per cluster; Na: number of alleles; Ho: observed heterozygosity; He: expected heterozygosity; Fis: inbreeding coefficient with 95% confidence interval. The “total” row contains the sum of N, the total number of alleles (Na), and the mean for the other indices.

| RPPs ($q \geq 80\%$) | N | Na | Ho | He | Fis |
| --- | --- | --- | --- | --- | --- |
| RPP1 | 134 | 128 | 0.684 | 0.691 | 0.005 [-0.052; 0.086] |
| RPP2 | 301 | 159 | 0.662 | 0.67 | 0.011 [-0.005; 0.034] |
| Total | 435 | 179 | 0.673 | 0.681 | 0.008 |

**Fig. 2:**
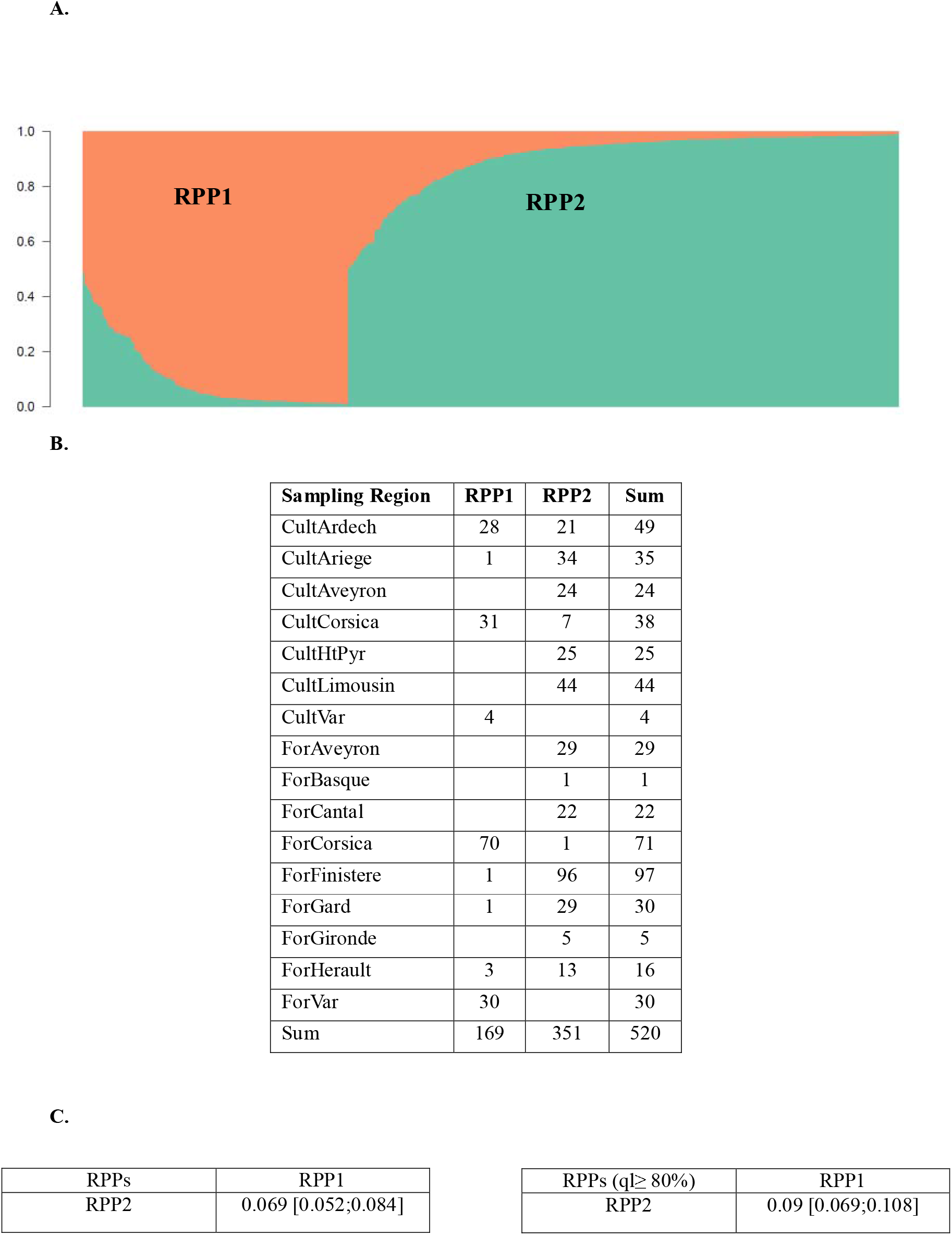
A: Classification of 520 European chestnut genotypes, in reconstructed panmictic populations (RPPs) when K =2, based on the *18Unik* data set. In orange, genotypes from south-east of France and Corsica (RPP1). In green, genotypes from other regions (RPP2). B: **Table of assignment of sampling regions in RPPs when K =2, based on the *18Unik* data set. Tables of pairwise Fst between RPPs using all individuals and strongly assigned individuals (ql ≥ 80%), when K =2, based on the *18Unik* data set.**

In terms of assignment probabilities of individuals to clusters, the “clusteredness” statistic was markedly higher for STRUCTURE and DAPC than for SNMF, for both *18Unik* at K=2 (0.809 and 0.909 versus 0.642) and *10Unik* at K=6 (0.732 and 0.922 versus 0.512).

## 4. DISCUSSION

### 4.1. Sampling

This work is the first comprehensive survey of genetic diversity and structure of *Castanea sativa* Mill. in France. As such, it fills the sampling gap in France for the benefit of future studies of chestnut structure in Europe. Our study benefited from two projects (the first author’s PhD and the FCBA project) which had different goals but whose sampling regions partially overlapped, and which used the same genotyping and allele scoring procedures. Combining these projects resulted in a large sampling effort to better assess the overall diversity of cultivated and forest chestnut in France, a crucial component of landscape genetics (Schwartz and McKelvey 2009).

### 4.2. Diversity indices

The levels of diversity in our sampling regions are comparable with those reported in other studies (Lusini et al. 2014; Mattioni et al. 2017; Mattioni et al. 2013; Skender et al. 2017), similarly, the mean numbers of alleles per locus are comparable with those obtained in other European regions (Lusini et al. 2014; Pereira-Lorenzo et al. 2017).

### 4.3. Null alleles frequencies

Some SSRs were known to often have a high null allele frequency, such as EmCs25 (Lusini et al. 2014) and CsCAT14, CsCAT2, CsCAT41B, QrZAG4 and CIO (Pereira-Lorenzo et al. 2017). In our data sets, EmCs38 had between 14 and 19 alleles, CIO had 6 and EmCs25 had 7. These loci had a high null allele frequency because of an excess of homozygotes.

### 4.4. Redundant diversity among sampling regions and no differentiation between chestnut types

The absence of significant genetic structure between forest and cultivated stands, and the high variance found within sampling regions, implies that each sampling region hosts substantial diversity, mostly shared with the other sampling regions. Such redundancy between sampling regions can be interpreted as the result of human exchanges (Bruneton-Governatori 1999; Conedera et al. 2016; Krebs et al. 2019; Pitte 1986).

Redundant genetic diversity in our sampling regions should ensure backup diversity, as long as information about landraces is shared among stakeholders in the different sampling regions. *In situ* sampling revealed that many landraces are multi-clonal. This source of diversity and hence of potential adaptation argues in favor of not reducing a landrace to one arbitrary clone. Even clones should be carefully evaluated, as morphological differences between individual trees attributed to the same clones were reported during our field trips, as has been the case in other species (Cipriani et al. 2010). All this is particularly interesting at a time when chestnut valuation tends to be based on heritage, with significance and quality marks based on local landraces (e.g., AOC Châtaigne d’Ardèche, AOC Farine de châtaigne Corse – Farina castagnina corsa, Label rouge Marron du Périgord). Some studies could provide authorities with arguments to justify certifying that landraces are “local”. On the other hand, even if a landrace has been cultivated for centuries in a particular place, this may also be the case elsewhere. Therefore, one might rightfully ask whether the quality of local chestnut comes from its locality. For crops like chestnut, usage and practices may be at least as important as genetics to give value to chestnut for growers and consumers (Dupré 2002, 2005; Martin et al. 2017).

### 4.5. A highly admixed genetic structure

All methods identified a clear genetic structure in all data sets, with at least two clusters. The criterion applied to the STRUCTURE results provided the most clear-cut signal from which to choose the number of clusters, we hence based our analysis on these results. Moreover, the results with less SSRs but more individuals (*10Unik* data set) are coherent with the results with more SSRs but less individuals (*18Unik* data set). Several articles showed limits to STRUCTURE (e.g., Kalinowski, 2011; Puechmaille 2016) or to the way it can be used (Wang, 2016), but a full comparison with the two other methods we tested based on simulations is out of the scope of this article. Still, it is important to note that, at the number of clusters determined by STRUCTURE (K=2 for *18Unik* and K=6 for *10Unik*), the two other methods, DAPC and SNMF, assigned individuals to clusters similarly to STRUCTURE. Moreover, even though the SNMF model is closer to the STRUCTURE model than the model behind DAPC (K-means), its assignments of individuals to clusters were less differentiated, as shown by the “clusteredness” statistic.

Characterizing genetic diversity (respectively structure) as high (strong) or low (weak) can be particularly risky as it must be in relative terms. The main finding here was the high admixture between the regions we sampled, both forest and cultivated. Like (Pereira-Lorenzo et al. 2019), we could not separate wild chestnut from cultivated ones. The genetic closeness was confirmed by the AMOVA and a Fct close to zero between wild and cultivated chestnut. This result was not completely unexpected given that chestnut is an outcrossing species and that gene flow between forest and cultivated stands is known to occur, together with changes in usage over time and in certain practices such as forests being used as a source of seedlings for rootstock, good quality fruits as a source of seedlings to plant forests, peasant woods in Limousin (personal communication), and “instant domestication” (Pereira-Lorenzo et al. 2019).

### 4.6. Future outlook of European genetic structure of chestnut

The genetic structure inferred from our samples did not necessarily match the sampling regions. This result was also expected for a continuously dispersed species affected by human management like European chestnut. Moreover, an admixed genetic structure was consistent with the known patterns of divergence and distribution of chestnut (Mattioni et al. 2017), combined with evidence from fossil pollen of several tree species suggesting that chestnut populations originating from Italy or the Balkans spread into the Iberian Peninsula from the north (Grivet and Petit 2003; Petit 2003). In the EU database (2017) and in Pereira-Lorenzo et al. (2019), « Luguesa » was classified with the Italian group of cultivars. In our analyses, it was found in the south-eastern cluster (Online resource 8: RPP1 in 10Unik data set with Spanish samples at K=2) with « Pais», « Puga » and « Raigona », which were originally classified in the Iberian group. The other Spanish cultivars were found in cluster n°2. This classification of Spanish samples in the two main clusters detected is not sufficient to conclude on the spontaneous spread of the species from either Spanish or Italian last glacial refugia (Krebs et al. 2019). An update of the EU database including more French individual would be needed to clarify the origin of French chestnut.

Before removal of hybrid individuals, the Basque sampling region was represented in the 10 (respectively 18) dataset by 119 (respectively 10) successfully genotyped non-redundant individuals. This high number of admixed individuals is an important feature of the actual chestnut forest there, resulting from the long history of interspecific hybridization in this region which extends on both sides of the border between Spain and France (Pereira-Lorenzo et al. 2017). It is further substantiated by the high prevalence of trees tolerant to ink disease, as found through artificial inoculation experiments within seedlots we have genotyped (Robin et al., in preparation).

### 4.7. Future outlook of SNP genotyping

The markers we used were selected after an extensive review of the literature (by us for the 13 SSR, and independently by Pereira-Lorenzo et al. 2018), and allele scoring was the subject of a recent optimization by Pereira-Lorenzo et al. (2018). Nevertheless, we faced the usual difficulties and drawbacks of microsatellites, i.e., errors and uncertainties in allele calling, difficulty in data comparison and transferability across labs and collaborators over time, and the huge amount of time needed to perform the analysis, as emphasized in previous studies (reviewed by Guichoux et al. 2011).

We consequently set up a small project to define nuclear SNPs, at least to check clear duplicates (in the case of good quality genotyping results) and putative duplicate (in the case of low quality results) among samples from variety repositories. In a few months, we re-genotyped about 500 samples with up to 160 SNPs and confirmed all suspected duplicates. A detailed description of this work will be the subject of a separate article.

## 5. CONCLUSION

In conclusion, this study revealed the genetic diversity and structure of French forest and cultivated chestnut across most of its range. We showed high diversity redundancy between sampling regions and a weak genetic structure. Based on external knowledge, the influence of human activity is the most probable explanation for this finding. Two main clusters were found, one in Corsica and the south-east of France. This confirms existing historical knowledge on land use changes, the movement of landraces, and « instant domestication » landraces. We believe our work provides useful information for conservation planning purposes and for cooperation between chestnut non-profit associations and groups of growers interested in landrace conservation and diffusion.

## Supporting information

Online resource 7

Online Resource 6

Online resource 5

Online resource 4

Online resource 3

Online resource 2

Online resource 1

Online resource 9

Online resource 8

## ACKNOWLEDGMENTS

This paper is part of the PhD of the first author who is grateful of her supervisors Laurent Hazard and Nathalie Couix of the INRAE, AGIR and Timothée Flutre, INRAE, AGAP and Le Moulon.

We would thank the local partners who introduced the first author to the chestnut, helped in sampling and shared their knowledge. In particular members of ACRC, Châtaigne des Pyrénées, Rénova, SPCV and Paysans du Rance.

We would like to thank the people who provided access to forest samples or sampling regions:

In Corsica: GRPTCMC and INRAE Corsica; In Limousin: Croqueurs de pommes du Limousin and the Parc Naturel Régional Perigord-Limousin; In Ardèche and Eastern Occitanie: Chambre départementale d’agriculture d’Ardèche (CDA Ardèche), Syndicat des producteurs de châtaigne d’Ardèche (SDCA) and chambre régionale d’agriculture d’Occitanie (CRA Occitanie). Other public and private landowners who granted us access to their forest stands in Brittany, Gard, Aveyron, Basque country and Cantal

We are very grateful to advisors and partners taking part to other aspects of our work on chestnut, Invenio, and CTIFL, INRA, Biogeco, INRA, Biologie du Fruit et Pathologie, CIF-Lourizan (Spain), and the members of the national workgroup on chestnut forestry set up by the National Center for Private Forest owners (CNPF) and particularly its chairman René Lempire as well as, Jean Lemaire and Sabine Girard.

We would also like to thank FCBA colleagues involved in sampling.

The authors thank the anonymous reviewers for critically reading the manuscript and suggesting substantial improvements.

## Funding

A grant of The Fondation de France and a regional research program for and on rural development (PSDR4-Occitanie, France) fund the PhD of the first author.

In 2016/2017, local partners (Rénova, ACRC and Châtaigne des Pyrénées) collected subsidies from the Conservatory of the Regional Biological Patrimony (CPBR) for the genotyping.

The FCBA project on forest samples was funded by the Conseil Régional de Nouvelle Aquitaine through the “sélection châtaignier à bois” project (2015-2016) contrat number 15007198-046 and grant “FEDER-FSA 2014-2020 Dossier 47010”, and “sélection châtaignier” project, grant n°16008302-043 and “FEDER-FSE 2014-2020 – AXE 1 n° 3296610”. The two FEDER grant parts are funds obtained from European Union by the Conseil Régional de Nouvelle Aquitaine. We thank the members of the FCBA external advisory board on Forestry and members of the National Center for private forest owners for its help to setup and manage the above-mentioned projects

The Xylobiotech platform is funded through the Equipe Xyloforest project (grant ANR-10-EQPX-16).

## Data availability

The datasets analysed during the current study are available in the data.inra repository at a private link https://data.inra.fr/privateurl.xhtml?token=8c03a83c-be4d-4984-972f-7808558b4539. The link will be published after the review process will be completed.

