## Supplementary material for "High admixture between forest an cultivated chestnut (*Castanea sativa* Mill.) in France": Online resource 7

ESM7: Detection of genetic clusters

### I. *18UNIK* data set

#### A. BIC of the *18Unik* data set

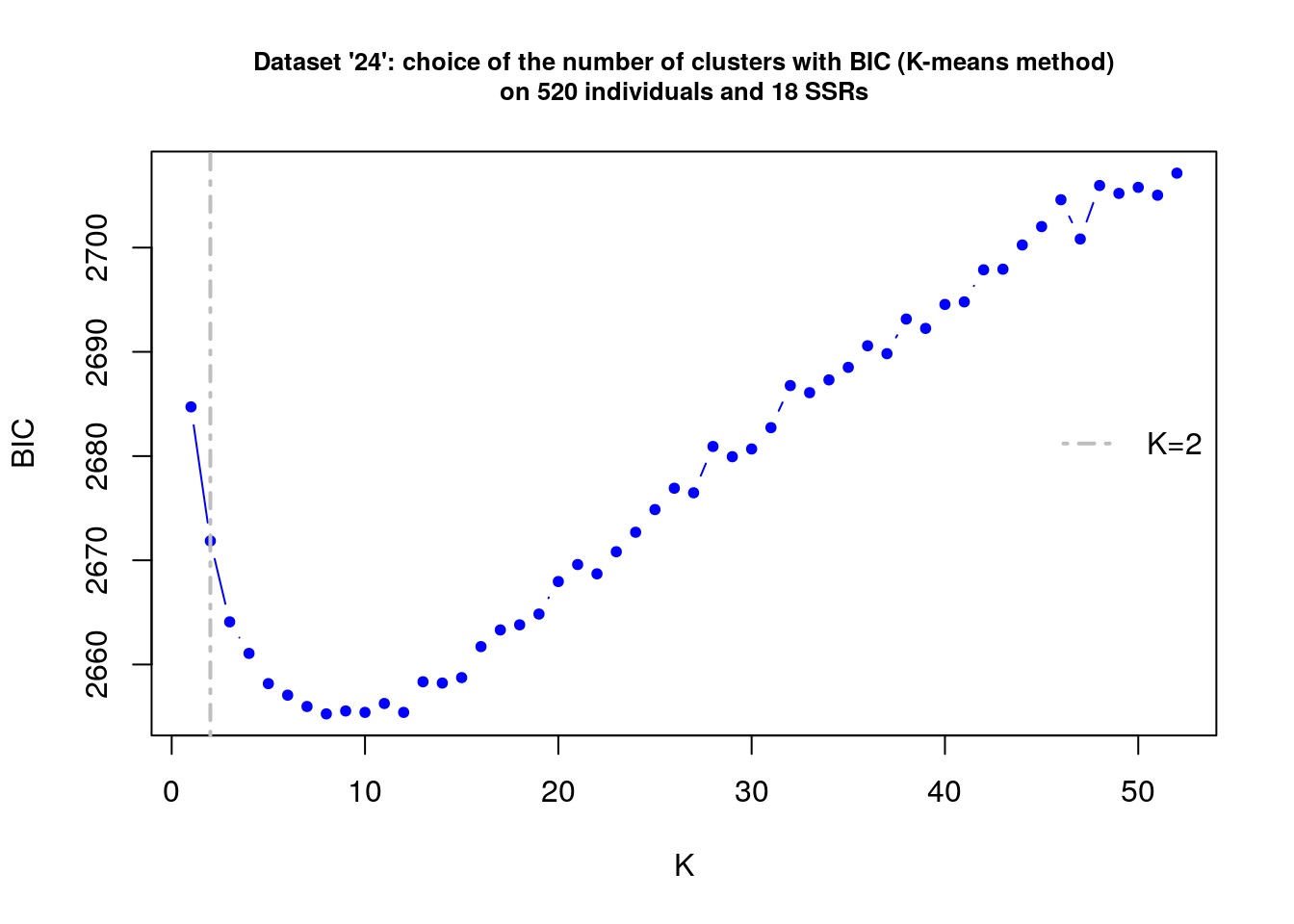

#### B. Cross-entropy criterion of the *18Unik* data set

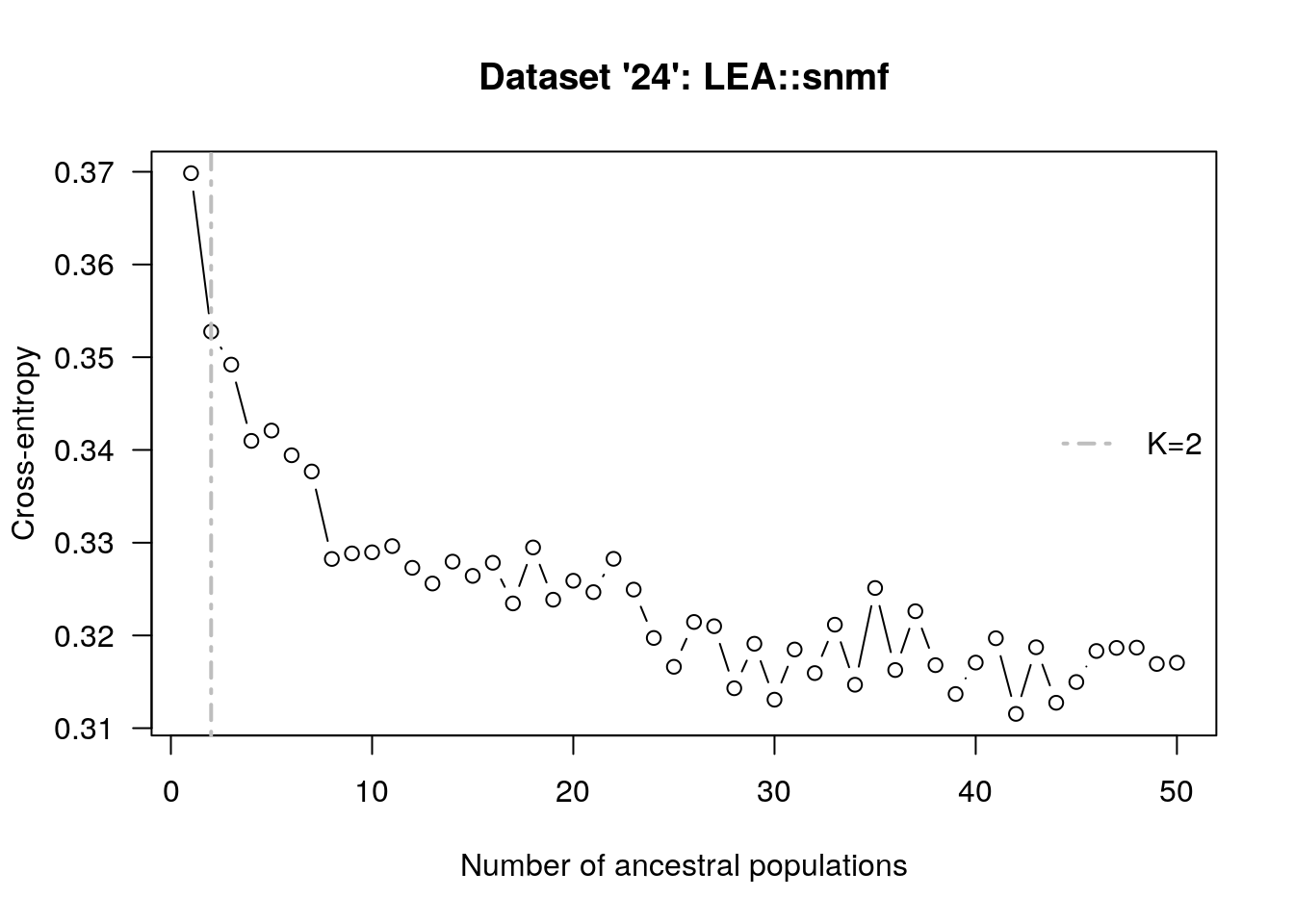

#### C. DeltaK of the *18Unik* data set

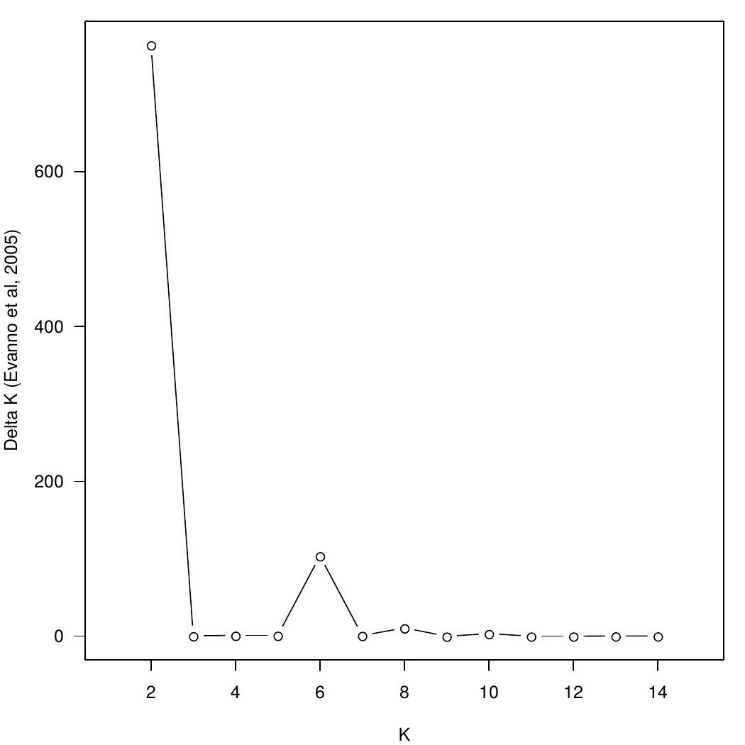

### II. *10Unik* data set

#### A. BIC of the *10Unik* data set

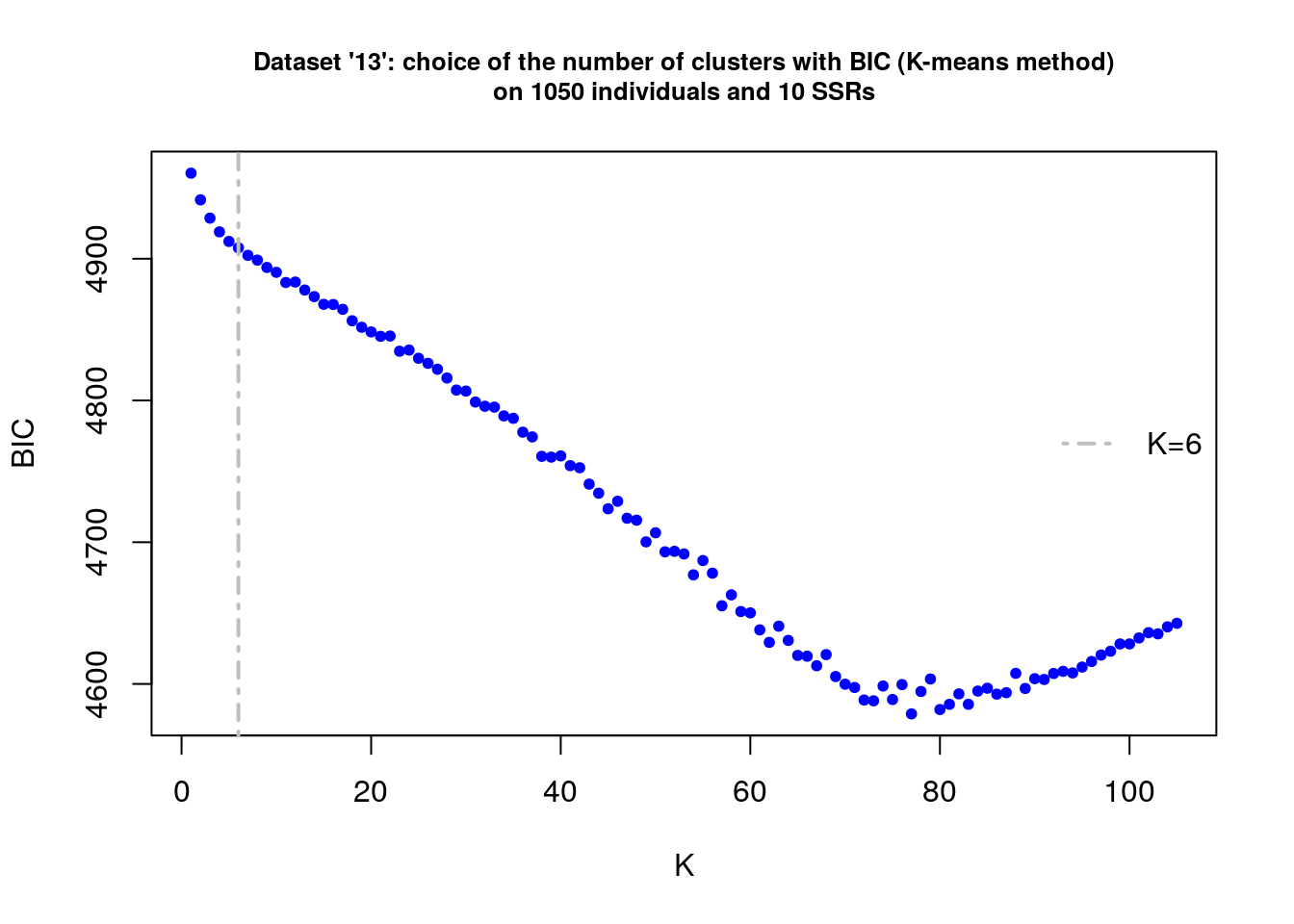

#### B. Cross-entropy criterion of the *10Unik* data set

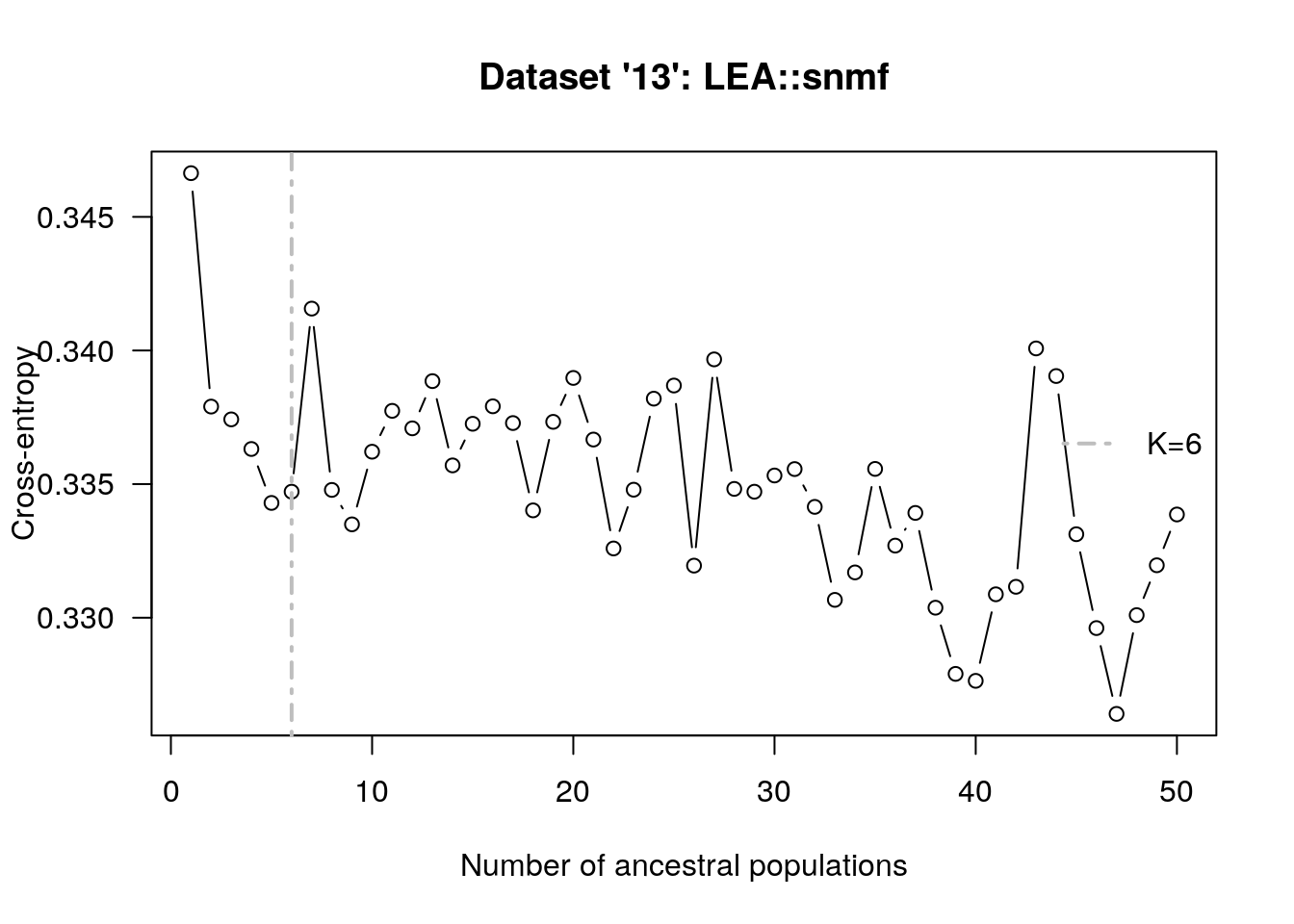

#### C. DeltaK of the *10Unik* data set

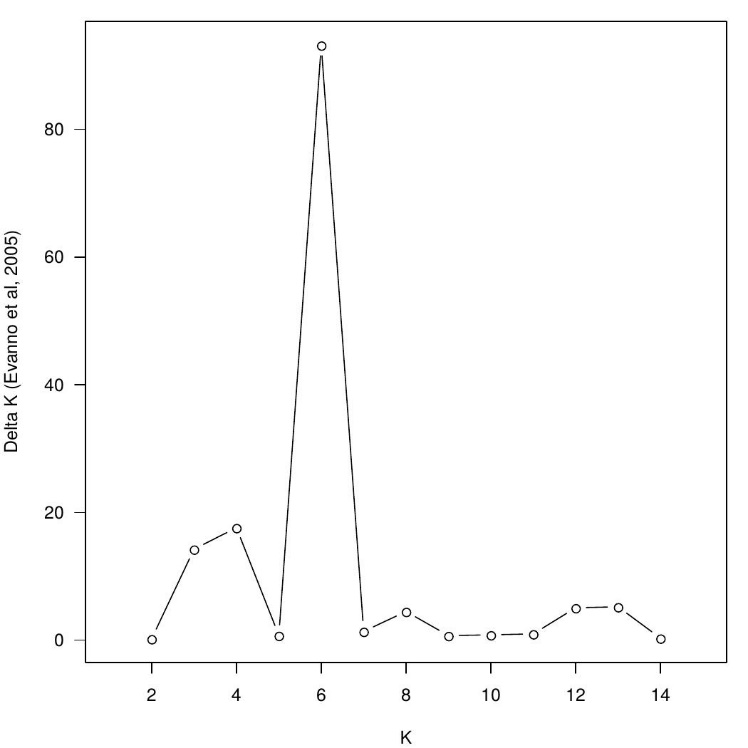

### III. *18Unik* data set with spanish samples

#### A. BIC of the *18Unik* data set with spanish samples

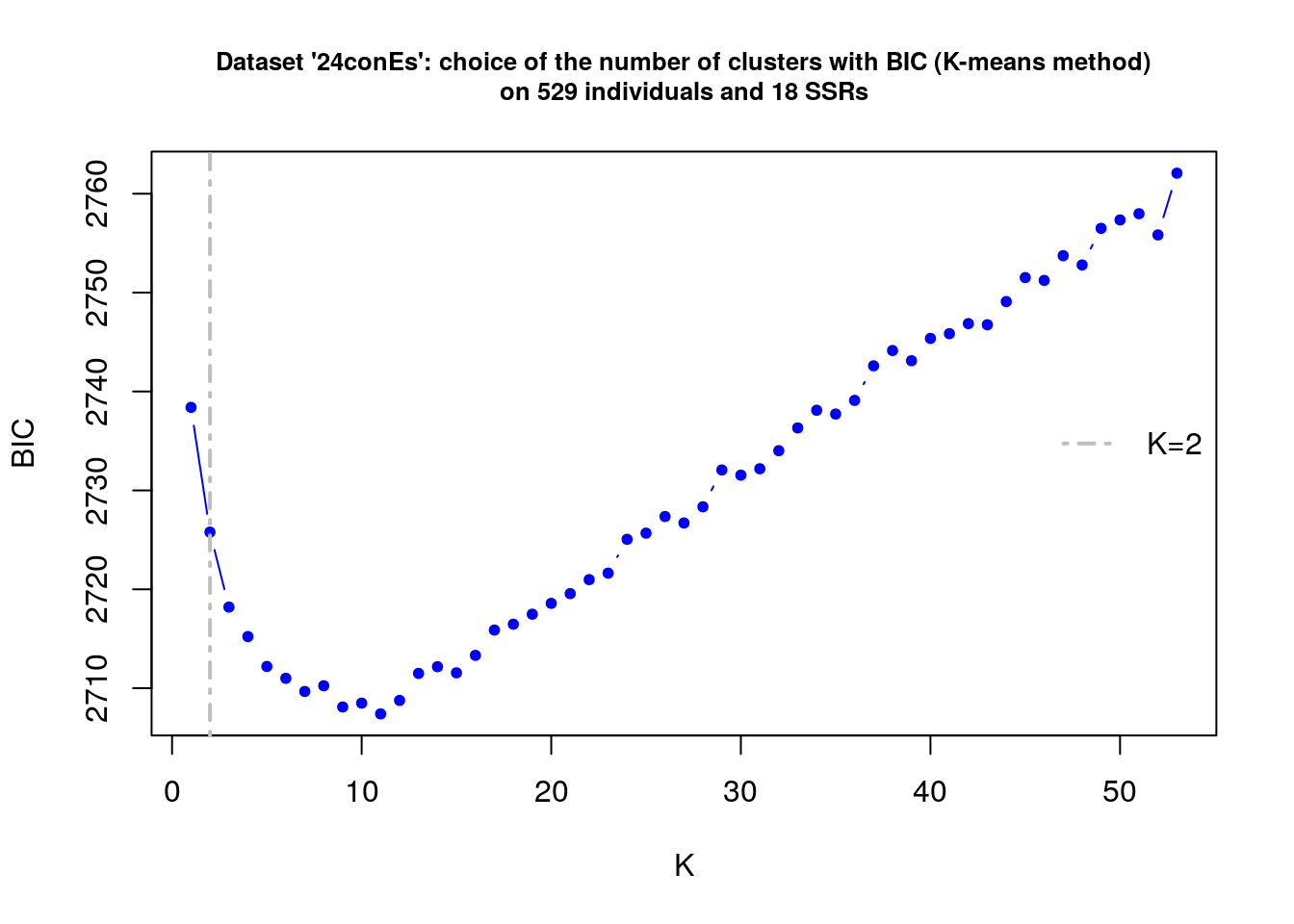

#### B. Cross-entropy criterion of the *18Unik* data set with spanish samples

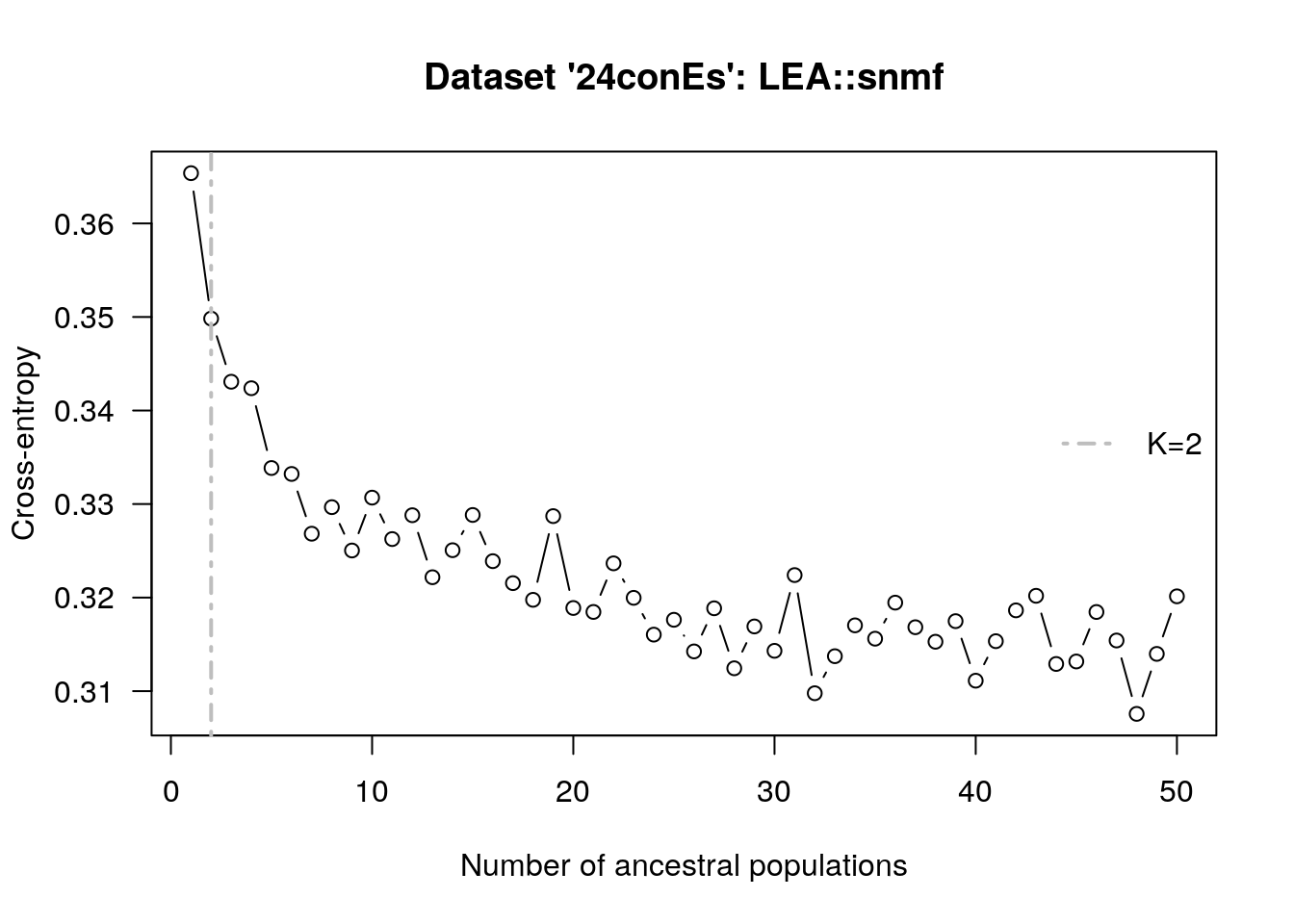

#### C. DeltaK of the *18Unik* data set with spanish samples

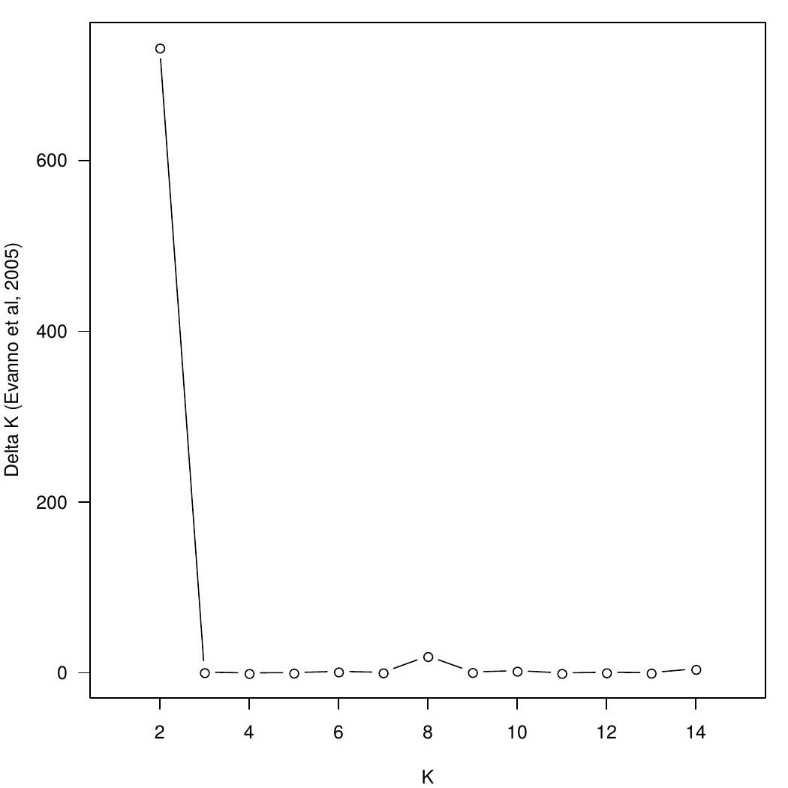

### IV. *10Unik* data set with spanish samples

#### A. BIC of of the *10Unik* data set with Spanish samples

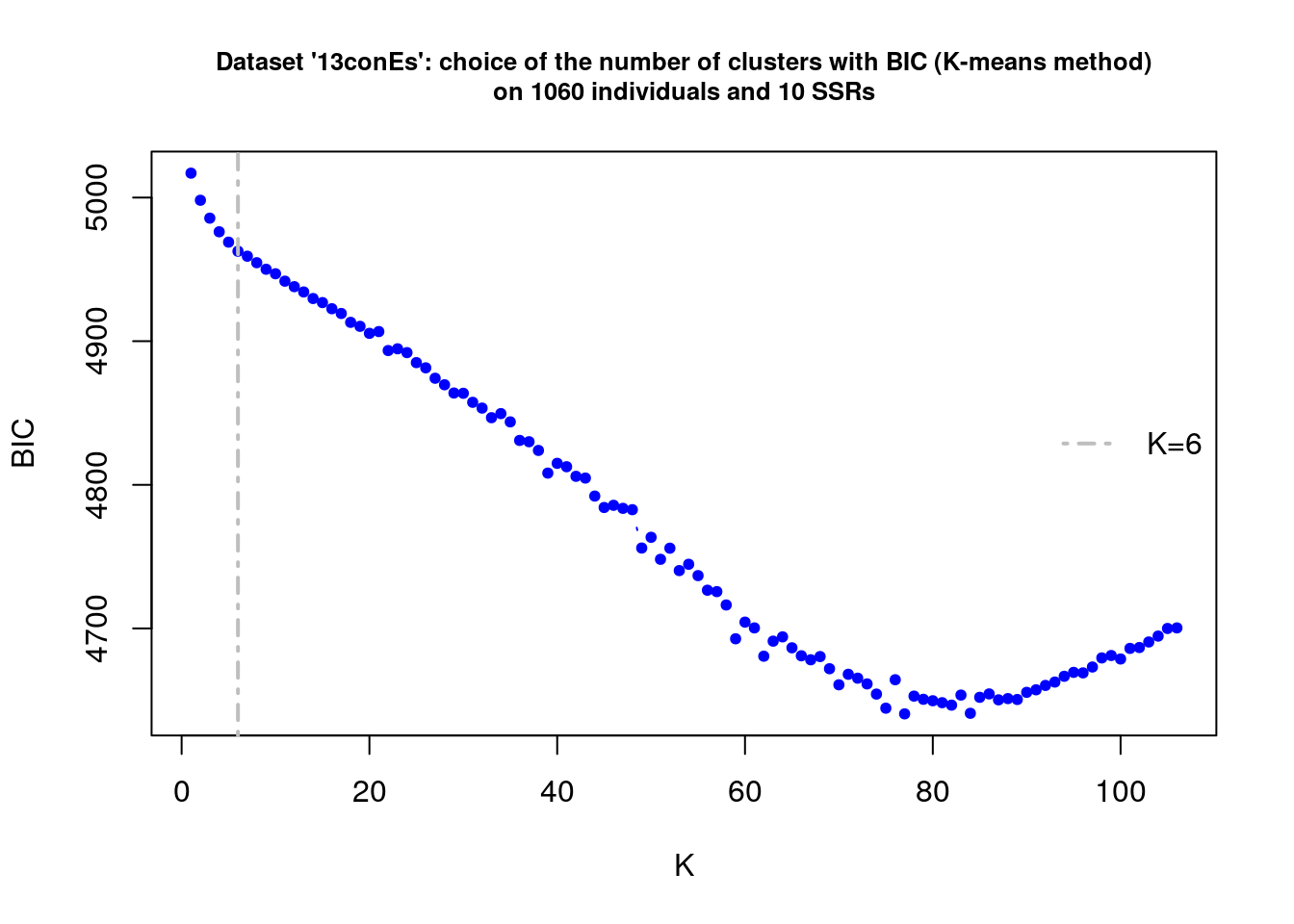

#### B. Cross-entropy criterion of the *10Unik* data set with Spanish samples

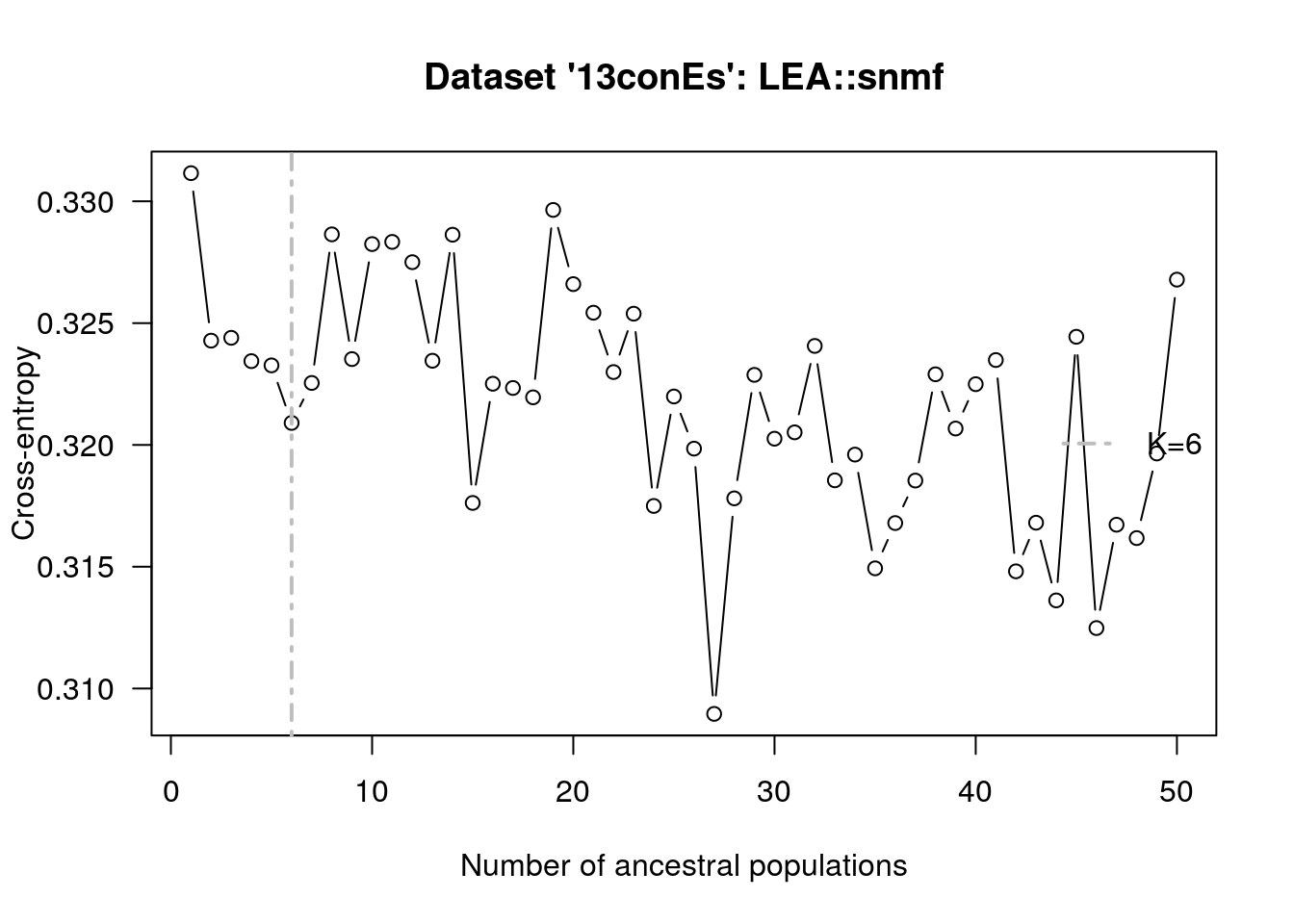

#### C. DeltaK of the *10Unik* data set with Spanish samples

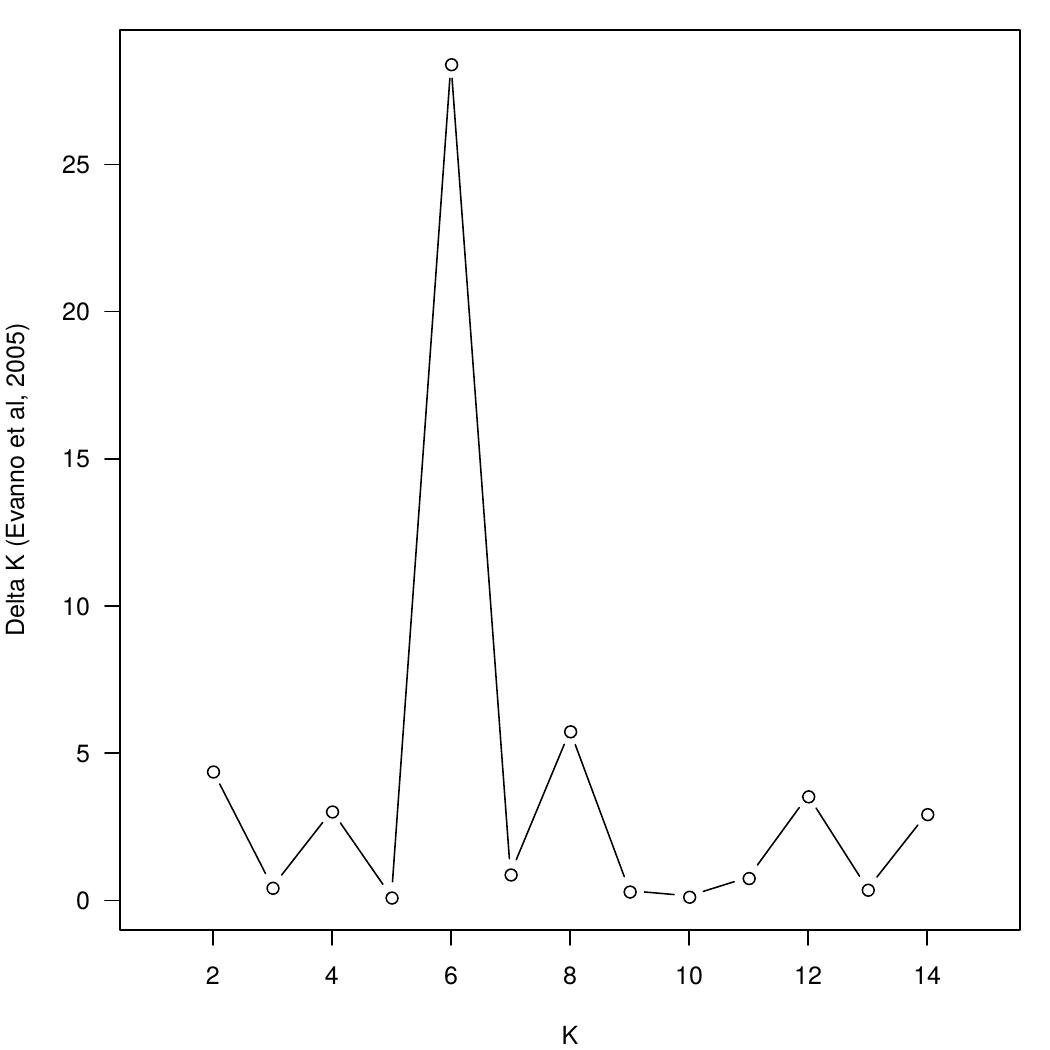
