## Supplementary material for "High admixture between forest an cultivated chestnut (*Castanea sativa* Mill.) in France": Online Resource 6

**ESM6: Hierarchical AMOVA and F-statistics**

I. **ESM6A: Hierarchical AMOVA and F-statistics for 17 French sampling regions genotyped at 10 SSRs without MLGs (*10Unik* data set)**

df: degrees of freedom, Alter: alternative hypothesis, 95% confidence intervals,*** p value ≤ 0.001.

| **Source of variation** | **df** | **Variance component** | **% of variation** | **p value** | **Alter** | **F statistic** |
| --- | --- | --- | --- | --- | --- | --- |
| **Among chestnut type** | 1 | -0.078 | -1.23 | 0.831 | greater | Fct -0.007 [-0.011 ; -0.002] |
| **Among sampling regions within chestnut types** | 15 | 0.799 | 12.58 | 0.001*** | greater | Fst 0.083 [0.07 ; 0.0.95] |
| **Within sampling regions** | 1033 | 5.629 | 88.64 | 0.001*** | less | Fis 0.015 [-0.009 ; 0.046] |
| **Total** | 1049 | 6.351 | 100.00 |  |  | Fit 0.09 [-0.06 ; 0.125] |

II. **ESM6B: Hierarchical AMOVA and F-statistics for six genetic clusters genotyped at 10 SSRs without MLGs (*10Unik* data set)**

df: degrees of freedom, Alter: alternative hypothesis, 95% confidence intervals,*** p value ≤ 0.001

| **Source of variation** | **df** | **Variance component** | **% of variation** | **p value** | **Alter** | **F statistic** |
| --- | --- | --- | --- | --- | --- | --- |
| **Among Clusters** | 5 | 0.729 | 11.3 | 0.001*** | greater | Fst 0.078 [0.064 ; 0.091] |
| **Within Clusters** | 1044 | 5.709 | 88.7 |  |  | Fis 0.022 [-0.005 ; 0.055] |
| **Total** | 1049 | 6.438 | 100.00 |  |  | Fit 0.098 [0.063 ; 0.136] |

III. **ESM6C: Hierarchical AMOVA and F-statistics for six genetic clusters genotyped of strongly assigned individuals (ql≥ 80%) at 10 SSRs without MLGs (*10Unik* data set)**

df: degrees of freedom, Alter: alternative hypothesis, 95% confidence intervals,*** p value ≤ 0.001

| **Source of variation** | **df** | **Variance component** | **% of variation** | **p value** | **Alter** | **F statistic** |
| --- | --- | --- | --- | --- | --- | --- |
| **Among Clusters** | 5 | 1.21 | 18.1 | 0.001*** | greater | Fst 0.125 [0.104 ; 0.144] |
| **Within Clusters** | 558 | 5.46 | 81.9 |  |  | Fis 0.002 [-0.03 ; 0.042] |
| **Total** | 563 | 6.67 | 100.00 |  |  | Fit 0.126 [0.085 ; 0.173] |
