## Supplementary material for "High admixture between forest an cultivated chestnut (*Castanea sativa* Mill.) in France": Online resource 5

**ESM5: Genetic diversity parameters per chestnut sampling region
genotyped at 10 SSRs without MLGs (10*Unik* data set)**

N: number of unique individuals genotyped per sampling region; Na: number of alleles; Ne: mean number of effective alleles; Ho: observed heterozygosity; He: expected heterozygosity; H: Shannon-Weiner diversity index; Ia: index of association; rbarD: standardized index of association; Fis: inbreeding coefficient, with 95% confidence interval (CI). Stars indicate significant *p* values at the 0.001 threshold. The “Total” row contains the sum for N, the total for Na and H, and the mean for the other indices. In bold are extrema (for Fis, CI excluding 0).

| **Sampling Regions** | **N** | **Na** | **Ne** | **Ho** | **He** | **H** | **Ia** | **rbarD** | **Fis** |
| --- | --- | --- | --- | --- | --- | --- | --- | --- | --- |
| **CultArdech** | 47 | 63 | 3.11 | 0.696 | 0.671 | 3.85 | 0.495* | 0.055* | -0.080 [-0.191;0.032] |
| **CultAriege** | 64 | 68 | 3.37 | 0.627 | 0.698 | 4.16 | 0.274* | 0.031* | 0.055 [-0.055;0.148] |
| **CultAveyron** | 70 | 67 | 3.17 | 0.7 | 0.68 | 4.25 | 0.397* | 0.044* | -0.061 [-0.163;0.016] |
| **CultCorsica** | 38 | 62 | 3.41 | 0.684 | 0.697 | 3.64 | 0.285 | 0.032 | -0.013 [-0.136;0.116] |
| **CultHtPyr** | 42 | 62 | 3.14 | 0.618 | 0.673 | 3.74 | 0.445* | 0.05* | -0.041 [-0.064;0.175] |
| **CultLimousin** | 59 | 65 | 3.30 | 0.725 | 0.691 | 4.08 | 0.535* | 0.06* | -0.067 [-0.171;0.012] |
| **CultVar** | 13 | 39 | 2.22 | 0.598 | 0.529 | 2.56 | 2.851* | 0.333* | -0.137 [-0.302;0.069] |
| **ForArdech** | 86 | 65 | 3.20 | 0.642 | 0.683 | 4.45 | 0.217* | 0.024* | 0.015 [-0.086;0.103] |
| **ForAveyron** | 140 | 58 | 2.75 | 0.633 | 0.635 | 4.94 | 0.237* | 0.026* | -0.034 [-0.136;0.024] |
| **ForBasque** | 24 | 44 | 2.42 | 0.614 | 0.575 | 3.18 | 0.415* | 0.046* | -0.072 [-0.174;0.005] |
| **ForCantal** | 22 | 47 | 2.54 | 0.636 | 0.593 | 3.09 | 0.272 | 0.030 | -0.089 [-0.186 ;0.027] |
| **ForCorsica** | 116 | 69 | 3.45 | 0.680 | 0.707 | 4.75 | 0.185* | 0.021* | -0.005 [-0.112 ;0.111] |
| **ForFinistere** | 248 | 87 | 3.68 | 0.719 | 0.727 | 5.51 | 0.083* | 0.009* | -0.034 [-0.107;0.033] |
| **ForGard** | 30 | 56 | 2.90 | 0.740 | 0.645 | 3.40 | 0.251 | 0.028 | -0.161 [-0.260 ;-0.079] |
| **ForGironde** | 5 | 41 | 3.60 | 0.580 | 0.650 | 1.61 | 1.043 | 0.124 | 0.205 [-0.022;0.386] |
| **ForHerault** | 16 | 49 | 3.20 | 0.719 | 0.666 | 2.77 | 0.286 | 0.032 | -0.081 [-0.193 ;0.141] |
| **ForVar** | 30 | 47 | 2.40 | 0.587 | 0.574 | 3.40 | 0.189 | 0.022 | -0.07 [-0.228;0.190] |
| **Total** | 1050 | 112 | 3.81 | 0.659 | 0.653 | 6.94 | 0.168* | 0.019* | 0.035 |
