## Supplementary material for "High admixture between forest an cultivated chestnut (*Castanea sativa* Mill.) in France": Online resource 4

**ESM4: Tests for Hardy-Weinberg equilibrium (HWE) on the *18Unik* and 10*Unik* data sets**

In order to test the HWE per sampling region, we first separated the sampling regions with the function seppop() from package adegenet, focusing on the analytical *p* values by setting B = 0. We then created a matrix that only contains *p* values, with sampling regions in columns and loci in rows. From this matrix we created the figures below which are heatmap showing significant departure from HWE. Note, that all loci shown in pink are loci suspected of not being in HWE with *p*≤0.05.

### ESM4A: Tests for HWE over all sampling regions (18Unik)

| **Locus** | **chi^2** | **df** | **Pr(chi^2>)** | **Pr.exact** |
| --- | --- | --- | --- | --- |
| **CsCAT14** | 36.14 | 10 | 0.000 | 0 |
| **CsCAT2** | 925.289 | 171 | 0.000 | 0 |
| **EmCS15** | 13.682 | 6 | 0.033 | 0.027 |
| **CsCAT16** | 110.958 | 45 | 0.000 | 0 |
| **CsCAT3** | 375.864 | 465 | 0.999 | 0.022 |
| **QpZAG36** | 28.894 | 10 | 0.001 | 0 |
| **CsCAT41B** | 717.913 | 55 | 0.000 | 0 |
| **QpZAG110** | 47.17 | 55 | 0.765 | 0.176 |
| **QrZAG4** | 0.554 | 1 | 0.457 | 0.597 |
| **QrZAG96** | 72.068 | 21 | 0.000 | 0 |
| **CsCAT1** | 125.739 | 55 | 0.000 | 0.102 |
| **CsCAT15** | 52.317 | 45 | 0.211 | 0.003 |
| **CsCAT6** | 467.523 | 120 | 0.000 | 0 |
| **CsCAT8** | 393.228 | 36 | 0.000 | 0 |
| **CsCAT17** | 97.665 | 45 | 0.000 | 0 |
| **RIC** | 3.585 | 6 | 0.733 | 0.653 |
| **OCI** | 24.222 | 10 | 0.007 | 0.007 |
| **OAL** | 59.518 | 36 | 0.008 | 0.011 |

### ESM4B: Heatmap of analytical p values from HWE tests per sampling region (18Unik)

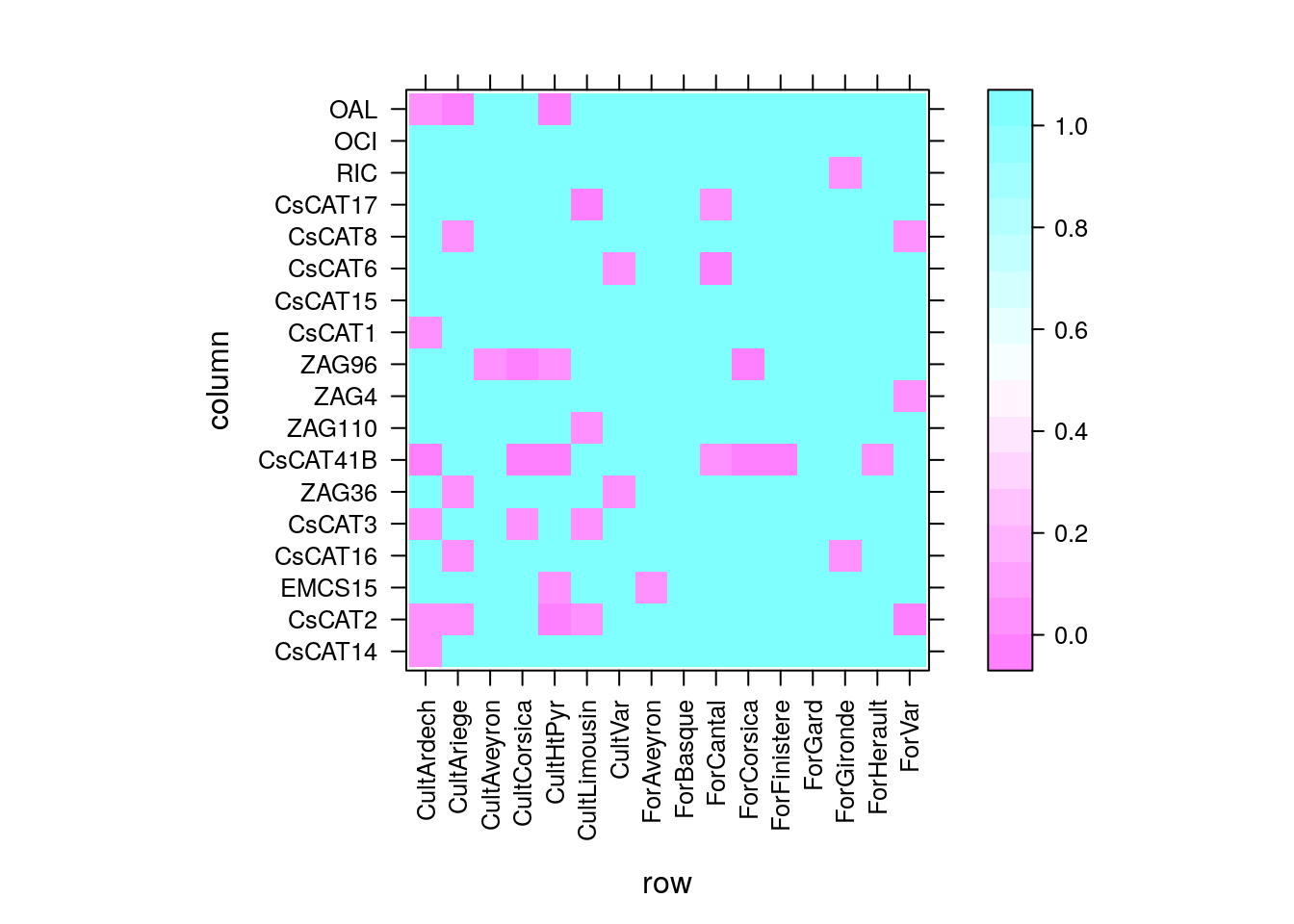

### ESM4C: Tests for HWE over all sampling regions (10Unik)

| **Locus** | **chi^2** | **df** | **Pr(chi^2>)** | **Pr.exact** |
| --- | --- | --- | --- | --- |
| **CsCAT14** | 60.4 | 15 | 0.000 | 0.000 |
| **CsCAT2** | 1716.4 | 190 | 0.000 | 0.000 |
| **EmCS15** | 31.7 | 6 | 0.000 | 0.000 |
| **EmCS2** | 25.1 | 3 | 0.000 | 0.000 |
| **CsCAT16** | 183.4 | 45 | 0.000 | 0.000 |
| **CsCAT3** | 544.1 | 528 | 0.305 | 0.003 |
| **QpZAG36** | 47.4 | 10 | 0.000 | 0.000 |
| **CsCAT41B** | 2044.5 | 78 | 0.000 | 0.000 |
| **QpZAG110** | 67.2 | 55 | 0.125 | 0.009 |
| **QrZAG96** | 107.0 | 21 | 0.000 | 0.000 |

### ESM4D: Heatmap of analytical p values from HWE tests per sampling region (10Unik)

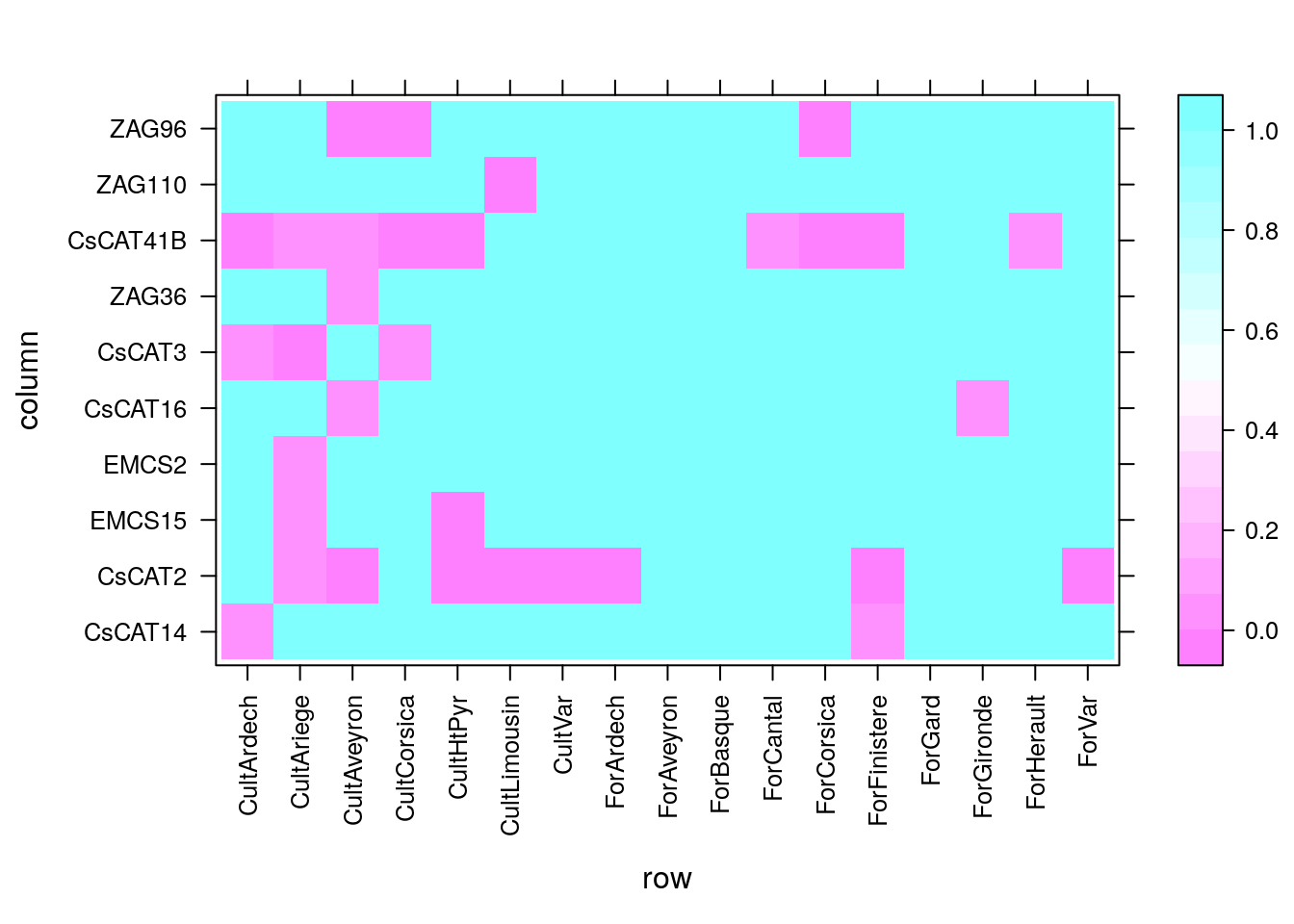
