## Supplementary material for "High admixture between forest an cultivated chestnut (*Castanea sativa* Mill.) in France": Online resource 3

**ESM3: Genetic diversity parameters per locus for the *Unik* data sets
(without MLGs)**

**ESM3A: Genetic diversity parameters per locus for the *18Unik*** **data set**

| **Locus** | **Range (pb)** | **Na** | **Ho** | **He** | **Fst** | **Fstp** | **Fis** | **Dest** |
| --- | --- | --- | --- | --- | --- | --- | --- | --- |
| **CsCAT14** | 134-161 | 5 | 0.65 | 0.69 | 0.083 | 0.088 | -0.002 | 0.17 |
| **CsCAT2** | 194-234 | 19 | 0.69 | 0.85 | 0.152 | 0.161 | 0.152 | 0.476 |
| **EmCs15** | 80-88 | 4 | 0.67 | 0.71 | 0.025 | 0.027 | -0.427 | 0.064 |
| **CsCAT16** | 128-158 | 10 | 0.73 | 0.75 | 0.058 | 0.062 | -0.06 | 0.145 |
| **CsCAT3** | 188-278 | 31 | 0.83 | 0.85 | 0.064 | 0.068 | -0.029 | 0.317 |
| **QpZAG36** | 207-219 | 5 | 0.72 | 0.72 | 0.091 | 0.097 | -0.126 | 0.180 |
| **CsCAT41B** | 210-233 | 11 | 0.58 | 0.77 | 0.158 | 0.167 | -0.224 | 0.340 |
| **QpZAG110** | 204-234 | 11 | 0.69 | 0.71 | 0.064 | 0.068 | -0.03 | 0.136 |
| **QrZAG4** | 109-113 | 2 | 0.18 | 0.17 | 0.159 | 0.167 | -0.275 | 0.043 |
| **QrZAG96** | 144-165 | 7 | 0.67 | 0.73 | 0.123 | 0.13 | -0.008 | 0.296 |
| **CsCAT1** | 177-223 | 11 | 0.7 | 0.7 | 0.092 | 0.097 | -0.131 | 0.196 |
| **CsCAT15** | 123-156 | 10 | 0.72 | 0.74 | 0.061 | 0.065 | -0.107 | 0.145 |
| **CsCAT6** | 157-200 | 16 | 0.79 | 0.86 | 0.125 | 0.132 | -0.106 | 0.469 |
| **CsCAT8** | 187-212 | 9 | 0.81 | 0.82 | 0.046 | 0.049 | -0.072 | 0.169 |
| **CsCAT17** | 131-160 | 10 | 0.81 | 0.82 | 0.091 | 0.096 | -0.103 | 0.288 |
| **RIC** | 119-127 | 4 | 0.67 | 0.67 | 0.063 | 0.067 | -0.123 | 0.120 |
| **OCI** | 144-158 | 5 | 0.69 | 0.7 | 0.114 | 0.120 | -0.061 | 0.231 |
| **OAL** | 297-332 | 9 | 0.52 | 0.54 | 0.107 | 0.114 | -0.097 | 0.109 |
| **Mean** | - | 9.94 | 0.673 | 0.711 | 0.093 | 0.099 | -0.102 | 0.216 |

**ESM3B: Genetic diversity parameters per locus for the 10*Unik*** **data set**

| **Locus** | **Range(pb)** | **Na** | **Ho** | **He** | **Fst** | **Fstp** | **Fis** | **Dest** |
| --- | --- | --- | --- | --- | --- | --- | --- | --- |
| **CsCAT14** | 134-161 | 6 | 0.62 | 0.67 | 0.084 | 0.088 | -0.038 | 0.163 |
| **CsCAT2** | 194-240 | 20 | 0.7 | 0.84 | 0.14 | 0.147 | 0.139 | 0.476 |
| **EmCs15** | 80-88 | 4 | 0.66 | 0.72 | 0.026 | 0.027 | -0.431 | 0.065 |
| **EmCs2** | 158-163 | 3 | 0.6 | 0.66 | 0.145 | 0.153 | 0.006 | 0.225 |
| **CsCAT16** | 128-158 | 10 | 0.72 | 0.77 | 0.080 | 0.085 | -0.008 | 0.19 |
| **CsCAT3** | 188-278 | 33 | 0.82 | 0.85 | 0.057 | 0.061 | -0.009 | 0.282 |
| **QpZAG36** | 207-219 | 5 | 0.68 | 0.7 | 0.08 | 0.084 | -0.115 | 0.161 |
| **CsCAT41B** | 210-237 | 13 | 0.61 | 0.74 | 0.137 | 0.144 | 0.169 | 0.316 |
| **QpZAG110** | 204-234 | 11 | 0.67 | 0.7 | 0.072 | 0.076 | -0.007 | 0.153 |
| **QrZAG96** | 144-165 | 7 | 0.67 | 0.72 | 0.08 | 0.085 | -0.02 | 0.201 |
| **Mean** | - | 11,2 | 0.675 | 0.737 | 0.09 | 0.095 | -0.031 | 0.223 |
