## Supplementary material for "High admixture between forest an cultivated chestnut (*Castanea sativa* Mill.) in France": Online resource 2

**ESM2: Plots of genotype accumulation curves for the *18Unik* and 10*Unik* data sets**

Genotype accumulation curves are useful for determining the minimum number of loci necessary to discriminate between individuals in a population. This was performed by randomly sampling loci without replacement and counting the number of multilocus genotypes (MLGs).

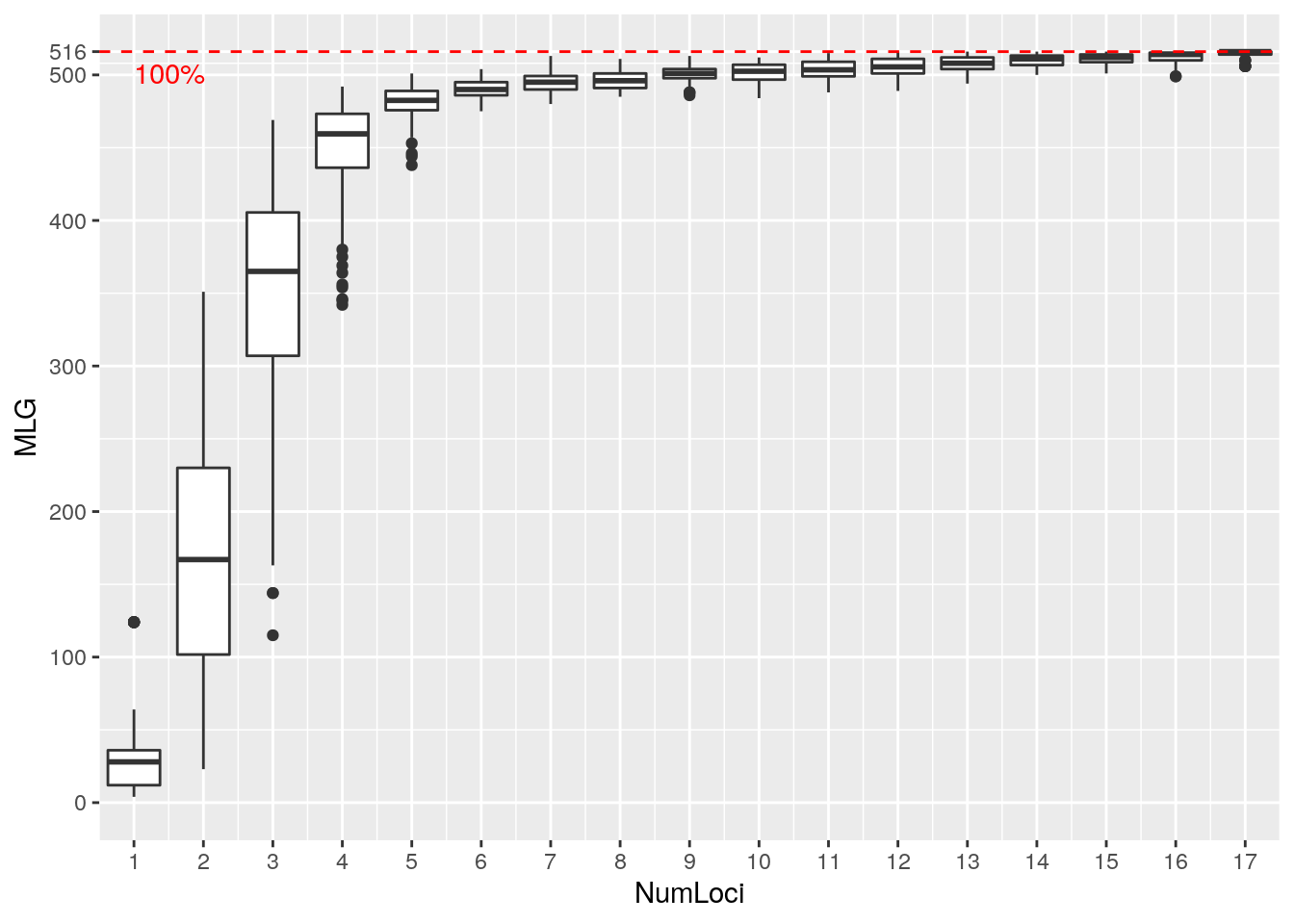

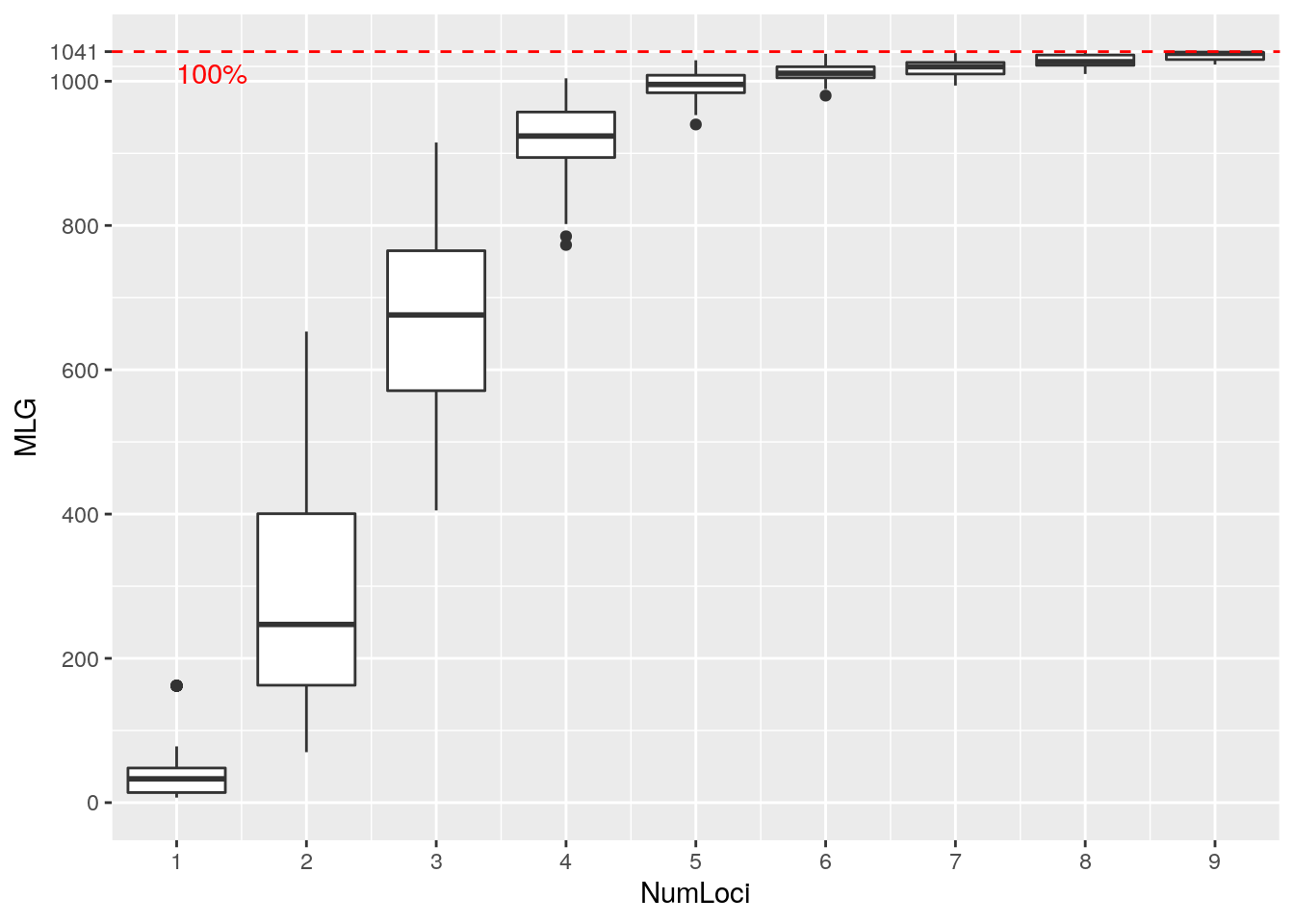
