## Supplementary material for "High admixture between forest an cultivated chestnut (*Castanea sativa* Mill.) in France": Online resource 1

**ESM1: Boxplots of null alleles according to Brookfield’s formula**

Any locus with a null allele frequency above a threshold of 10% was discarded: EmCs38, CIO and EmCs25 from the data set with 24 SSRs (first plot); EmCs38 from the data set with 13 SSRs (second plot).

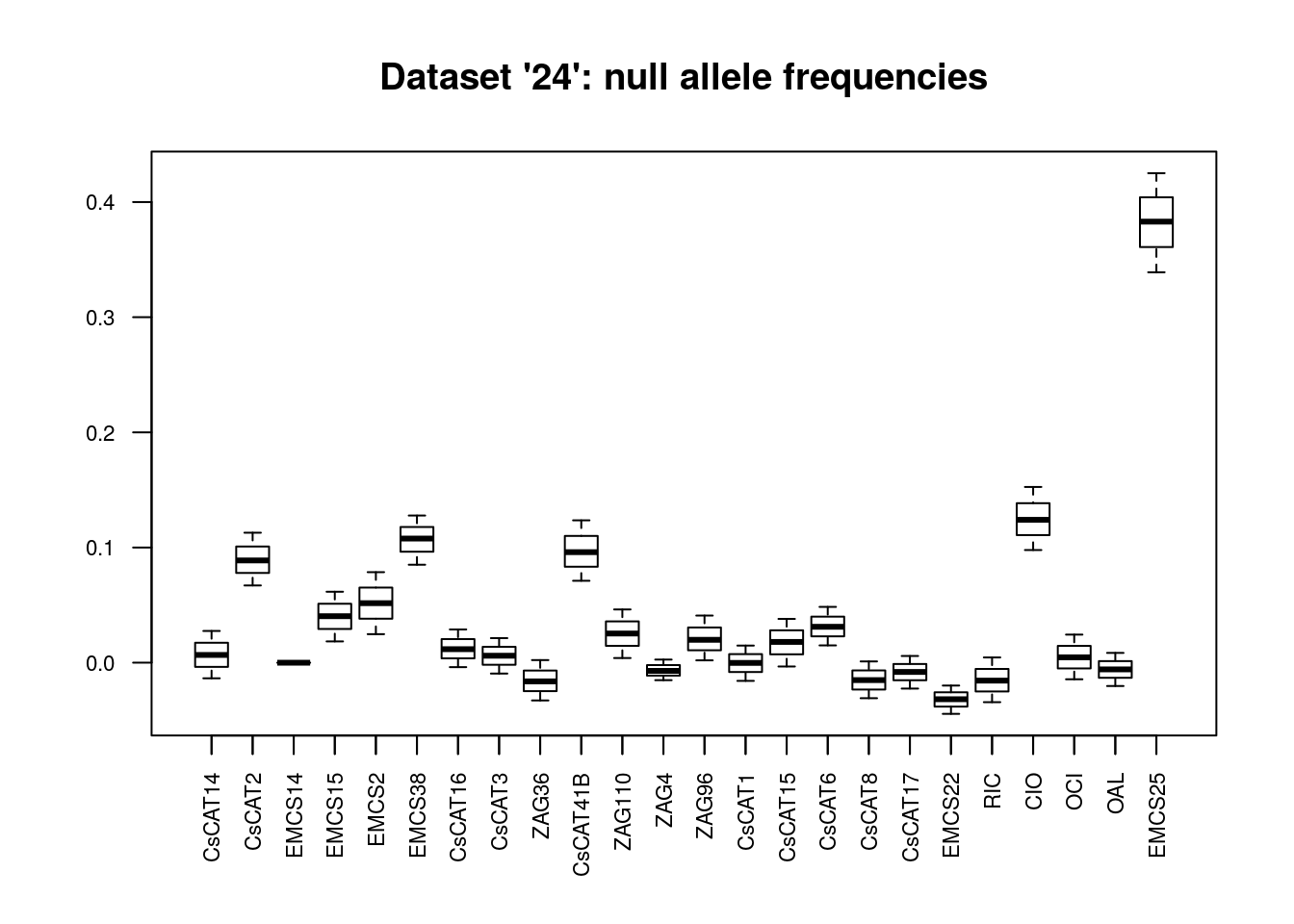

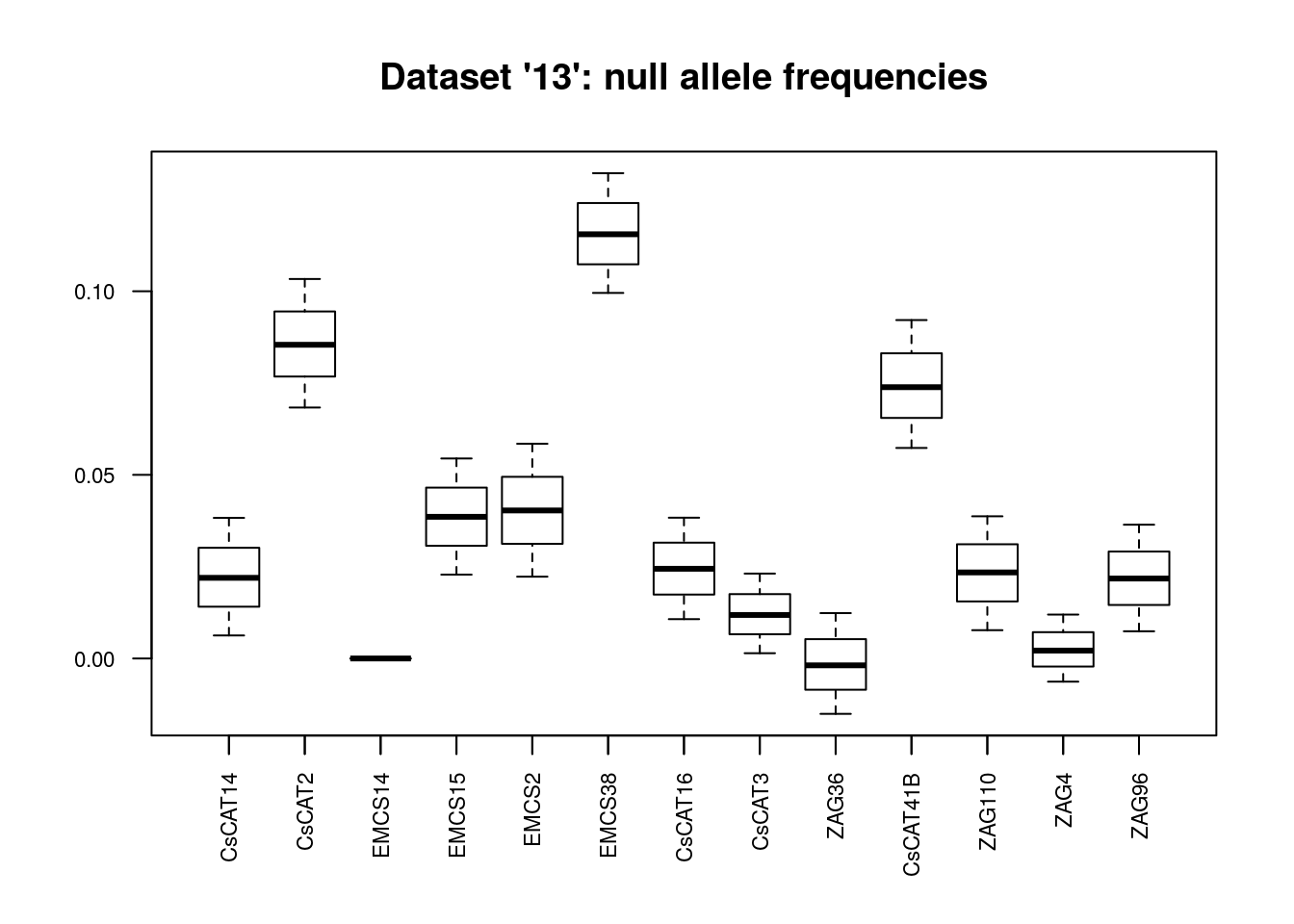
