## Supplementary material for "High admixture between forest an cultivated chestnut (*Castanea sativa* Mill.) in France": Online resource 9

**ESM9**: **Within-cluster genetic variability at 10 loci without MLGs, on strongly assigned individuals (ql≥ 80%)**

N: number of unique individuals genotyped per cluster; Na: number of alleles; Ho: observed heterozygosity; He: expected heterozygosity; Fis: inbreeding coefficient with 95% confidence interval. The “total” row contains the sum of N, the total number of alleles (Na), and the mean for the other indices.

| **RPPs**  **(ql≥ 80%)** | **N** | **Na** | **Ho** | **He** | **Fis** |
| --- | --- | --- | --- | --- | --- |
| **RPP1a \| Cluster 3** | 42 | 46 | 0.592 | 0.545 | -0.081 [-0.245;0.18] |
| **RPP2a \| Cluster 4** | 127 | 72 | 0.733 | 0.711 | -0.029 [-0.06;0.011] |
| **RPP2b\| Cluster 5** | 119 | 53 | 0.627 | 0.613 | -0.016 [-0.042;0.007] |
| **RPP2d \| Cluster 6** | 74 | 66 | 0.625 | 0.681 | 0.088 [0.026;0.153] |
| **RPP2c \| Cluster 2** | 96 | 63 | 0.676 | 0.675 | 0.003 [-0.03,0.032] |
| **RPP1b \| Cluster 1** | 106 | 70 | 0.686 | 0.702 | 0.024 [-0.067;0.141] |
| **Total** | 564 | 112 | 0.657 | 0.654 | -0.002 |
