## Supplementary material for "High admixture between forest an cultivated chestnut (*Castanea sativa* Mill.) in France": Online resource 8

ESM8

### I.  Classification of 529 European chestnut genotypes, in reconstructed panmictic populations (RPPs) when K=2, based on *18Unik* data set with spanish samples.

In green, French genotypes admixed (RPP1). In orange, genotypes from south-east of France (RPP2). Spanish genotypes are allocated in both RPPs.

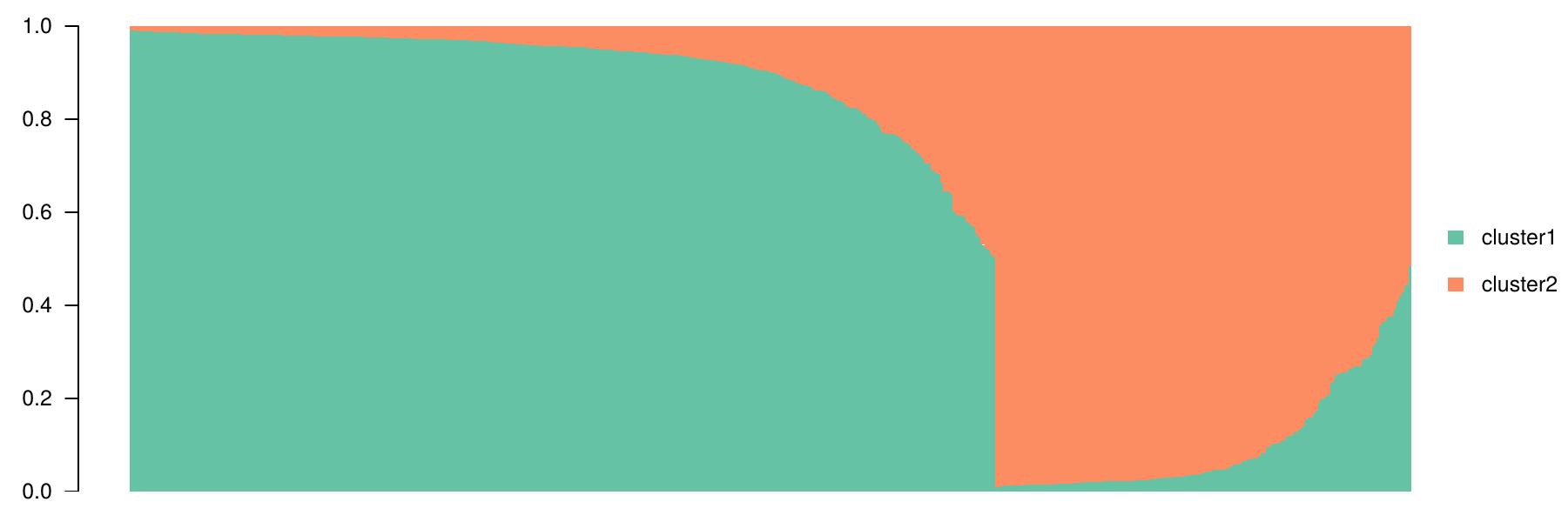

**RPP2**

**RPP1**

**Table of assignments of sampling regions in reconstructed panmictic populations (RPPs)**

| **Sampling Region** | **RPP1** | **RPP2** | **Sum** |
| --- | --- | --- | --- |
| **CultArdech** | 30 | 19 | 49 |
| **CultAriege** | 1 | 34 | 35 |
| **CultAveyron** |  | 24 | 24 |
| **CultCorsica** | 32 | 6 | 38 |
| **CultHtPyr** |  | 25 | 25 |
| **CultLimousin** |  | 44 | 44 |
| **CultSpain** | 6 | 3 | 9 |
| **CultVar** | 4 |  | 4 |
| **ForAveyron** |  | 29 | 29 |
| **ForBasque** |  | 1 | 1 |
| **ForCantal** |  | 22 | 22 |
| **ForCorsica** | 70 | 1 | 71 |
| **ForFinistere** | 1 | 96 | 97 |
| **ForGard** | 1 | 29 | 30 |
| **ForGironde** |  | 5 | 5 |
| **ForHerault** | 4 | 12 | 16 |
| **ForVar** | 30 |  | 30 |
| **Sum** | 179 | 350 | 529 |

**Table of posterior probabilities of assignment of Spanish samples in reconstructed panmictic populations (RPPs)**

| Samples | Name | RPP1 | RPP2 |
| --- | --- | --- | --- |
| ref002 | **Chamberga1** | 0.316 | **0.684** |
| ref003 | **Negral** | 0.289 | **0.711** |
| ref004 | **Pais** | 0.151 | **0.849** |
| ref005 | **Parede** | **0.69** | 0.31 |
| ref006 | **Porteliña** | **0.628** | 0.372 |
| ref007 | **Puga** | 0.273 | **0.727** |
| ref008 | **Raigona1** | 0.336 | **0.664** |
| ref009 | **Rapada** | 0.399 | **0.601** |
| ref010 | **Serodia** | **0.634** | 0.366 |

**Tables of pairwise Fst between RPPs using all individuals and strongly assigned individuals (ql≥ 80%)**

| RPPs | RPP1 |
| --- | --- |
| RPP2 | 0.065 [0.049 ; 0.08] |

| RPPs (ql≥ 80%) | RPP1 |
| --- | --- |
| RPP2 | 0.089 [0.069 ; 0.11] |

### II. Classification of 1050 European chestnut genotypes. in reconstructed panmictic populations (RPPs) when K=2 and K=6

## K=2

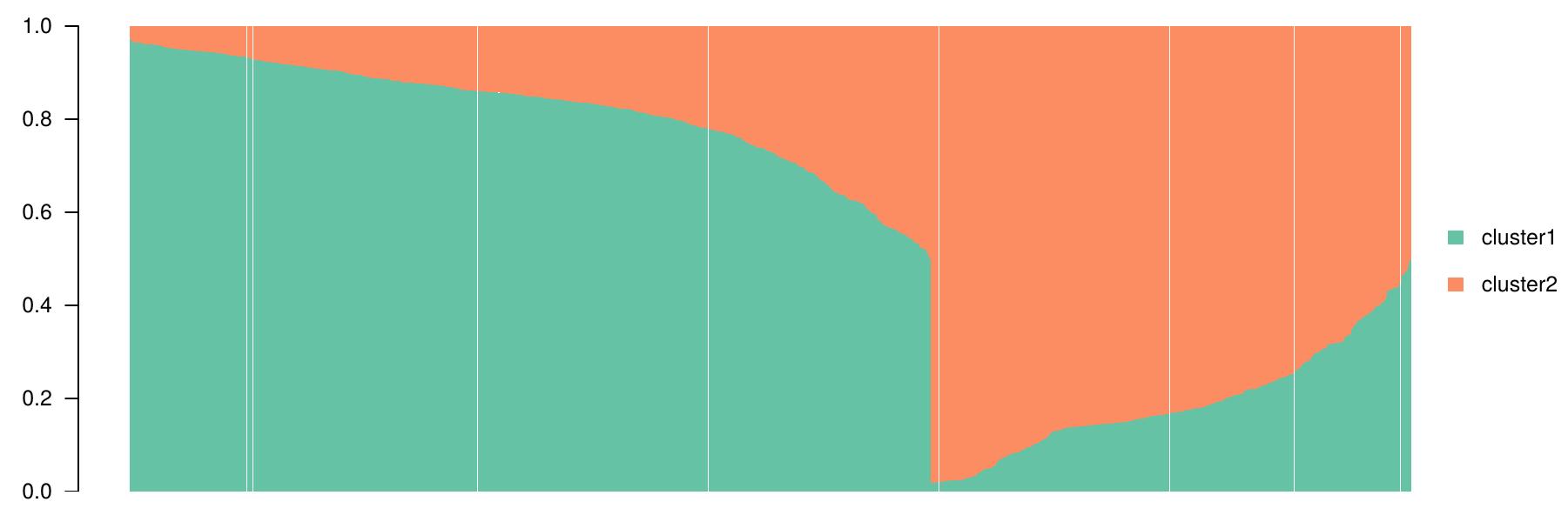

**RPP1**

**RPP2**

**Table of assignments of sampling regions in reconstructed panmictic populations (RPPs) when K=2**

| **Sampling Regions** | **cluster1 \|RPP2** | **cluster2 \| RPP1** | **Sum** |
| --- | --- | --- | --- |
| **CultArdech** | 6 | 41 | 47 |
| **CultAriege** | 41 | 23 | 64 |
| **CultAveyron** | 51 | 19 | 70 |
| **CultCorsica** | 1 | 37 | 38 |
| **CultHtPyr** | 29 | 13 | 42 |
| **CultLimousin** | 59 |  | 59 |
| **CultVar** | 1 | 12 | 13 |
| **ForArdech** | 40 | 46 | 86 |
| **ForAveyron** | 135 | 5 | 140 |
| **ForBasque** | 22 | 2 | 24 |
| **ForCantal** | 22 |  | 22 |
| **ForCorsica** | 1 | 115 | 116 |
| **ForFinistere** | 232 | 16 | 248 |
| **ForGard** | 10 | 20 | 30 |
| **ForGironde** | 1 | 4 | 5 |
| **ForHerault** | 5 | 11 | 16 |
| **ForVar** |  | 30 | 30 |
| **Sum** | 656 | 394 | 1050 |

**Tables of pairwise Fst between RPPs using all individuals and strongly assigned individuals (ql≥ 80%)**

| RPPs | RPP1 |
| --- | --- |
| RPP2 | 0.005 [0.029 ; 0.073] |

| RPPs (ql≥ 80%) | RPP1 |
| --- | --- |
| RPP2 | 0.088 [0.054 ; 0.123] |

## K=6

- In green. main genotypes are from Corsica (Cluster 1, RPP1b).
- In orange. main genotypes are from many sampling regions (Cluster 2, RPP2c)
- In blue. main genotypes are from Var and Ardech (Cluster 3, RPP1a).
- In pink. main genotypes are from West of France (Cluster 4, RPP2a).
- In clear green. genotypes from Aveyron (Cluster 5, RPP2b)
- In yellow. main genotypes are from Pyrenees (Cluster 6, RPP2d)

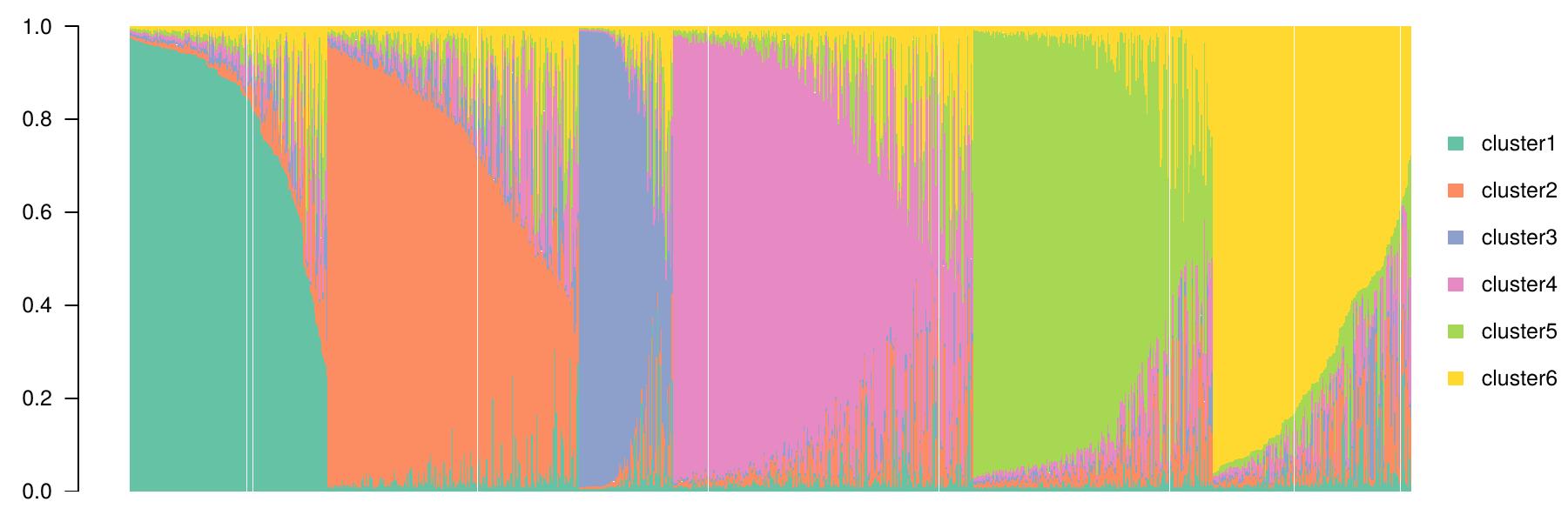

**RPP2d**

**RPP2b**

**RPP2a**

**RPP1a**

**RPP2c**

**RPP1b**

**Table of assignement of sampling regions in reconstructed panmictic populations (RPPs) when K=6**

| **Sampling Regions** | **RPP1b \| Cluster 1** | **RPP2c \| Cluster 2** | **RPP1a\| Cluster 3** | **RPP2a \| Cluster 4** | **RPP2b\|**  **Cluster 5** | **RPP2d \| Cluster 6** | **Sum** |
| --- | --- | --- | --- | --- | --- | --- | --- |
| **CultArdech** | 2 | 15 | 22 | 1 | 3 | 4 | 47 |
| **CultAriege** | 3 | 4 | 2 | 1 | 6 | 48 | 64 |
| **CultAveyron** |  | 19 | 5 | 14 | 24 | 8 | 70 |
| **CultCorsica** | 34 | 1 |  |  |  | 3 | 38 |
| **CultHtPyr** |  | 3 |  | 1 | 1 | 37 | 42 |
| **CultLimousin** |  | 3 |  | 43 | 4 | 9 | 59 |
| **CultVar** |  |  | 12 | 1 |  |  | 13 |
| **ForArdech** | 2 | 75 | 4 | 4 | 1 |  | 86 |
| **ForAveyron** |  | 11 |  | 2 | 123 | 4 | 140 |
| **ForBasque** | 2 | 2 |  | 1 |  | 19 | 24 |
| **ForCantal** |  | 1 |  | 1 | 17 | 3 | 22 |
| **ForCorsica** | 110 | 4 |  |  | 1 | 1 | 116 |
| **ForFinistere** | 6 | 39 | 3 | 173 | 13 | 14 | 248 |
| **ForGard** |  | 26 |  | 1 | 3 |  | 30 |
| **ForGironde** | 2 |  |  | 1 |  | 2 | 5 |
| **ForHerault** | 1 | 2 |  | 2 |  | 11 | 16 |
| **ForVar** |  | 1 | 29 |  |  |  | 30 |
| **Sum** | 162 | 206 | 77 | 246 | 196 | 163 | 1050 |

**Tables of pairwise Fst between RPPs using all individuals and strongly assigned individuals (ql≥ 80%)**

| **RPPs** | **RPP2c \| cluster2** | **RPP1a \| cluster3** | **RPP2a \| cluster4** | **RPP2b \| cluster5** | **RPP2d \| cluster6** |
| --- | --- | --- | --- | --- | --- |
| **RPP1b \| cluster1** | 0.062 [0.045;0.081] | 0.124 [0.071;0.193] | 0.077 [0.049;0.114] | 0.107 [0.082;0.13] |  |
| **RPP2c \| cluster2** |  | 0.103 [0.071;0.137] | 0.059 [0.035;0.085] | 0.075 [0.047;0.106] | 0.053 [0.031;0.081] |
| **RPP1a \| cluster3** |  |  | 0.153 [0.122;0.187] | 0.165 [0.110;0.214] | 0.142 [0.108;0.177] |
| **RPP2a \| cluster4** |  |  |  | 0.058 [0.033;0.082] | 0.037 [0.022;0.052] |
| **RPP2b \| cluster5** |  |  |  |  | 0.047 [0.026;0.070] |

| **RPPs**  **(ql≥ 80%)** | **RPP2c \| cluster2** | **RPP1a \| cluster3** | **RPP2a \| cluster4** | **RPP2b \| cluster5** | **RPP2d \| cluster6** |
| --- | --- | --- | --- | --- | --- |
| **RPP1b \| cluster1** | 0.099 [0.073;0.129] | 0.198 [0.115;0.286] | 0.101 [0.066;0.143] | 0.156 [0.116;0.192] | 0.122 [0.082;0.162] |
| **RPP2c \| cluster2** |  | 0.158 [0.105;0.207] | 0.096 [0.060;0.131] | 0.117 [0.074;0.164] | 0.097 [0.061;0.136] |
| **RPP1a \| cluster3** |  |  | 0.231 [0.18;0.286] | 0.245 [0.163;0.316] | 0.229 [0.177;0.282] |
| **RPP2a \| cluster4** |  |  |  | 0.088 [0.050;0.123] | 0.073 [0.048;0.095] |
| **RPP2b \| cluster5** |  |  |  |  | 0.079 [0.045;0.117] |

### III. Classification of 1060 European chestnut genotypes. in reconstructed panmictic populations (RPPs) when K=2 and K=6 with Spanish samples

## K=2

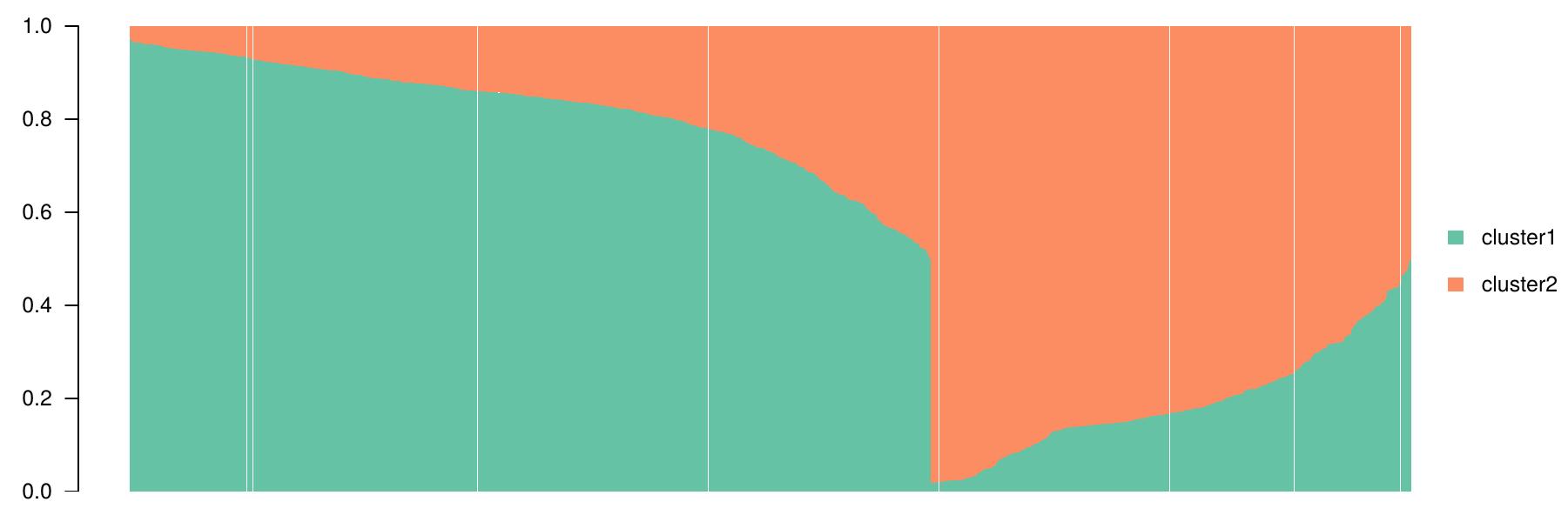

**RPP1**

**RPP2**

**Table of assignments of sampling regions in reconstructed panmictic populations (RPPs) when K=2**

| **Sampling Regions** | **cluster1 \|RPP2** | **cluster2 \| RPP1** | **Sum** |
| --- | --- | --- | --- |
| **CultArdech** | 6 | 41 | 47 |
| **CultAriege** | 42 | 22 | 64 |
| **CultAveyron** | 51 | 19 | 70 |
| **CultCorsica** | 1 | 37 | 38 |
| **CultHtPyr** | 30 | 12 | 42 |
| **CultLimousin** | 59 |  | 59 |
| **CultVar** | 1 | 12 | 13 |
| **CultSpain** | 3 | 7 | 10 |
| **ForArdech** | 39 | 47 | 86 |
| **ForAveyron** | 134 | 6 | 140 |
| **ForBasque** | 20 | 4 | 24 |
| **ForCantal** | 22 |  | 22 |
| **ForCorsica** | 1 | 115 | 116 |
| **ForFinistere** | 230 | 18 | 248 |
| **ForGard** | 10 | 20 | 30 |
| **ForGironde** | 1 | 4 | 5 |
| **ForHerault** | 5 | 11 | 16 |
| **ForVar** |  | 30 | 30 |
| **Sum** | 655 | 405 | 1060 |

**Tables of pairwise Fst between RPPs using all individuals and strongly assigned individuals (ql ≥ 80%)**

| RPPs | RPP1 |
| --- | --- |
| RPP2 | 0.005 [0.029 ; 0.072] |

| RPPs strongly assigned | RPP1 |
| --- | --- |
| RPP2 | 0.089 [0.054 ; 0.124] |

**Table of posterior probabilities of assignment of Spanish samples**

| **Samples** | **Name** | **Cluster1** | **Cluster2** |
| --- | --- | --- | --- |
| ref001 | **Luguesa** | 0.058 | **0.942** |
| ref002 | **Chamberga1** | 0.6703 | 0.3297 |
| ref003 | **Negral** | 0.4955 | 0.5045 |
| ref004 | **Pais** | 0.1407 | **0.8593** |
| ref005 | **Parede** | **0.8618** | 0.1382 |
| ref006 | **Porteliña** | 0.4929 | 0.5071 |
| ref007 | **Puga** | 0.1974 | **0.8026** |
| ref008 | **Raigona1** | 0.1474 | **0.8526** |
| ref009 | **Rapada** | 0.2068 | 0.7932 |
| ref010 | **Serodia** | 0.6479 | 0.3521 |

## B. K=6

- In green. main genotypes are from Corsica (Cluster 1, RPP1b).
- In orange. main genotypes are from many sampling regions (Cluster 2, RPP2c)
- In blue. main genotypes are from Var and Ardech (Cluster 3, RPP1a).
- In pink. main genotypes are from north west of France (Cluster 4, RPP2a).
- In clear green. genotypes from Aveyron (Cluster 5, RPP2b)
- In yellow. main genotypes are from Pyrenees (Cluster 6, RPP2d)

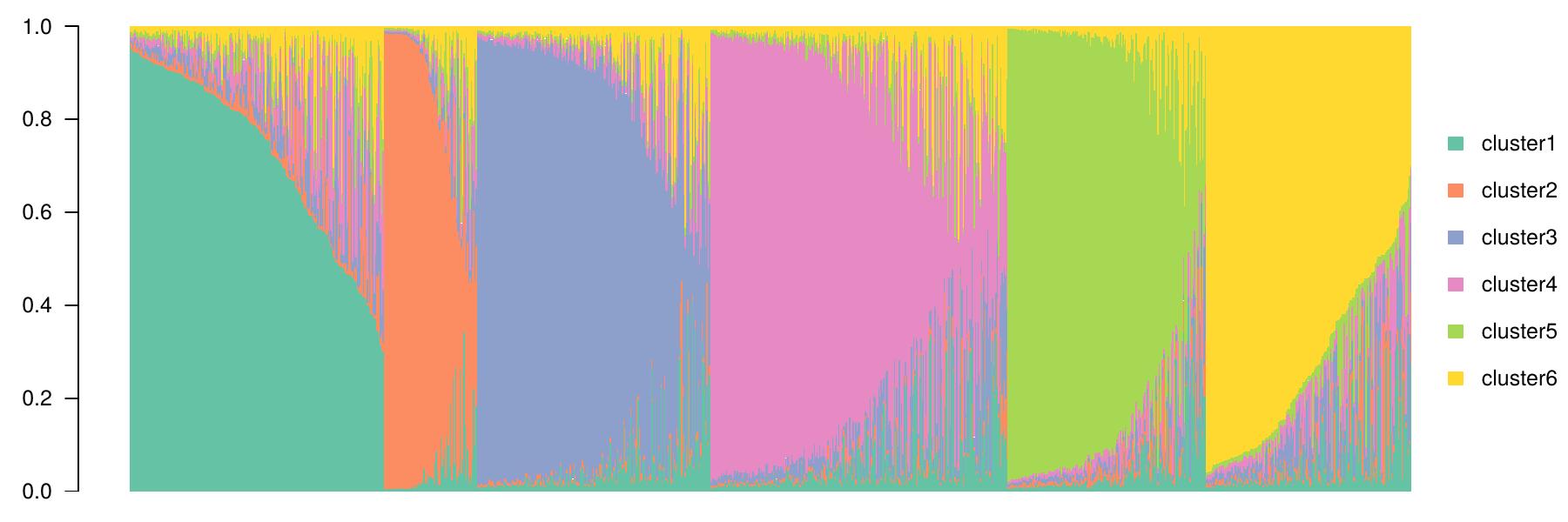

**RPP2d**

**RPP1b**

**RPP2a**

**RPP2b**

**RPP1a**

**RPP2c**

**Table of assignement of sampling regions in reconstructed panmictic populations (RPPs)**

| **Sampling Regions** | **RPP2c \| Cluster 1** | **RPP1a \| Cluster 2** | **RPP2b\| Cluster 3** | **RPP2a \| Cluster 4** | **RPP1b\|**  **Cluster 5** | **RPP2d \| Cluster 6** | **Sum** |
| --- | --- | --- | --- | --- | --- | --- | --- |
| **CultArdech** | 15 | 22 | 3 | 1 | 2 | 4 | 47 |
| **CultAriege** | 4 | 2 | 6 | 1 | 3 | 48 | 64 |
| **CultAveyron** | 19 | 5 | 21 | 18 |  | 7 | 70 |
| **CultCorsica** | 1 |  |  |  | 34 | 3 | 38 |
| **CultHtPyr** | 3 |  | 1 | 1 |  | 37 | 42 |
| **CultLimousin** | 4 |  | 4 | 42 |  | 9 | 59 |
| **CultVar** |  | 12 |  | 1 |  |  | 13 |
| **CultSpain** | 1 |  |  | 4 | 2 | 3 | 10 |
| **ForArdech** | 77 | 4 | 1 | 2 | 2 |  | 86 |
| **ForAveyron** | 11 |  | 123 | 2 |  | 4 | 140 |
| **ForBasque** | 2 |  |  | 1 | 2 | 19 | 24 |
| **ForCantal** | 1 |  | 17 | 1 |  | 3 | 22 |
| **ForCorsica** | 4 |  | 1 |  | 110 | 1 | 116 |
| **ForFinistere** | 39 | 3 | 13 | 168 | 6 | 19 | 248 |
| **ForGard** | 26 |  | 3 | 1 |  |  | 30 |
| **ForGironde** |  |  |  | 1 | 2 | 2 | 5 |
| **ForHerault** | 2 |  |  | 2 | 1 | 11 | 16 |
| **ForVar** | 1 | 29 |  |  |  |  | 30 |
| **Sum** | 210 | 77 | 193 | 246 | 164 | 170 | 1060 |

**Tables of pairwise Fst between RPPs using all individuals and strongly assigned individuals (ql ≥ 80%)**

| **RPPs** | **cluster2** | **cluster3** | **cluster4** | **cluster5** | **cluster6** |
| --- | --- | --- | --- | --- | --- |
| **cluster1** | 0.103 [0.072;0.135] | 0.075 [0.049;0.104] | 0.060 [0.036;0.085] | 0.061 [0.045;0.079] | 0.050[0.030;0.076] |
| **cluster2** |  | 0.166 [0.110;0.214] | 0.153 [0.121;0.185] | 0.123 [0.072;0.19] | 0.140 [0.105;0.172] |
| **cluster3** |  |  | 0.059 [0.035;0.085] | 0.107 [0.082;0.13] | 0.048 [0.027;0.072] |
| **cluster4** |  |  |  | 0.077 [0.048;0.112] | 0.037 [0.023;0.053] |
| **cluster5** |  |  |  |  | 0.071 [0.046;0.096] |

| **RPPs**  **(ql≥ 80%)** | **cluster2** | **cluster3** | **cluster4** | **cluster5** | **cluster6** |
| --- | --- | --- | --- | --- | --- |
| **cluster1** | 0.157 [0.104;0.207] | 0.118 [0.076;0.162] | 0.097 [0.06;0.135] | 0.100 [0.075;0.128] | 0.096 [0.063;0.135] |
| **cluster2** |  | 0.245 [0.163;0.316] | 0.232 [0.181;0.286] | 0.199 [0.119;0.287] | 0.231 [0.181;0.281] |
| **cluster3** |  |  | 0.088 [0.053;0.123] | 0.157 [0.116;0.194] | 0.082 [0.045;0.120] |
| **cluster4** |  |  |  | 0.101 [0.066;0.143] | 0.071 [0.047;0.093] |
| **cluster5** |  |  |  |  | 0.121 [0.080;0.161] |

**Table of posterior probabilities of assignment of Spanish samples**

| **Samples** | **Name** | **Cluster1** | **Cluster2** | **Cluster3** | **Cluster4** | **Cluster5** | **Cluster6** |
| --- | --- | --- | --- | --- | --- | --- | --- |
| ref001 | **Luguesa** | 0.3809 | 0.2989 | 0.007 | 0.007 | 0.275 | 0.0312 |
| ref002 | **Chamberga1** | 0.0223 | 0.0052 | 0.0101 | 0.0414 | 0.036 | **0.8849** |
| ref003 | **Negral** | 0.0216 | 0.0103 | 0.0192 | 0.4945 | 0.4108 | 0.0436 |
| ref004 | **Pais** | 0.0289 | 0.0525 | 0.0137 | 0.0334 | **0.7942** | 0.0773 |
| ref005 | **Parede** | 0.0161 | 0.0149 | 0.0769 | **0.7942** | 0.0648 | 0.0332 |
| ref006 | **Porteliña** | 0.0367 | 0.0101 | 0.0088 | 0.5012 | 0.1989 | 0.2442 |
| ref007 | **Puga** | 0.0093 | 0.0617 | 0.01 | 0.0267 | 0.2627 | 0.6296 |
| ref008 | **Raigona1** | 0.0213 | 0.314 | 0.0083 | 0.0317 | 0.1454 | 0.4792 |
| ref009 | **Rapada** | 0.0353 | 0.049 | 0.048 | 0.0543 | 0.5119 | 0.3014 |
| ref010 | **Serodia** | 0.0132 | 0.0062 | 0.0131 | 0.6748 | 0.2157 | 0.077 |
